# Hematopoietic stem and progenitor cells integrate *Bacteroides*-derived innate immune signals to promote gut tissue repair

**DOI:** 10.1101/2021.10.05.463122

**Authors:** Yoshikazu Hayashi, Maiko Sezaki, Gaku Nakato, Subinoy Biswas, Tatsuya Morishima, Md Fakruddin, Jieun Moon, Soyeon Ahn, Pilhan Kim, Yuji Miyamoto, Hideo Baba, Shinji Fukuda, Hitoshi Takizawa

## Abstract

Bone marrow (BM)-resident hematopoietic stem and progenitor cells (HSPCs) are often activated by bacterial insults to replenish the host hemato-immune system, but how they integrate the associated tissue damage signals to initiate distal tissue repair is largely unknown. Here, we showed that acute gut inflammation expands HSPCs in the BM through GM-CSFR activation, and directs them to inflamed mesenteric lymph nodes for further differentiation into myeloid cells specialized in gut tissue repair. We also identified that this process is exclusively mediated by *Bacteroides*, a commensal gram-negative bacteria, that activates innate immune signaling. In contrast, chronic gut inflammation reduces HSC potential for hematopoietic reconstitution and immune response against infection. Similarly, microbial signals contribute to aging-associated HSPC expansion. These findings establish a cross-organ communication that promotes tissue regeneration, but if sustained, impairs tissue homeostasis that may be relevant to aging and chronic disorders.

**Summary:** The infiltrating microbiota *Bacteroides* upon acute colitis directed MPP migration from the BM to the MLN for their subsequent expansion and differentiation into tissue-repairing Ly6C^+^/G^+^ cells, whereas chronic colitis impairs HSC functionality similarly as aging.

## Introduction

Adult hematopoietic stem cells (HSCs) slowly but continuously self-renew and differentiate into all blood lineages through intermediate progenitor cells, i.e., multipotent progenitors (MPPs) in order to sustain lifelong hematopoiesis (Orkin and Zon, 2008). This stem cell property is tightly controlled and preserved in the bone marrow (BM) microenvironment which provides pivotal factors for HSC maintenance. Although HSCs divide very infrequently and are mostly dormant at steady-state, they can be activated to enhance mature blood cell production upon demand, such as in cases of hematopoietic challenges like infection or inflammation (Takizawa et al., 2012). During bacterial or viral infections, HSCs are activated by pro-inflammatory cytokines that reach the BM, such as interferons and macrophage-colony stimulating factor (M-CSF) secreted from immune cells that recognize infection in peripheral tissues (Baldridge et al., 2010; Burberry et al., 2014; Essers et al., 2009; Mossadegh-Keller et al., 2013; Sato et al., 2009). HSCs can also sense infection directly through pattern recognition receptors (PRRs) such as Toll-like receptors (TLRs) that recognize pathogen-associated molecular patterns (PAMPs) of bacteria or viruses, and drive myelopoiesis at the expense of lymphopoiesis in order to replenish innate immune cells consumed at the site of infection. The response of HSCs to infection is overall beneficial, leading to their increased proliferation, differentiation or migration (Liu et al., 2015; Massberg et al., 2007; Megias et al., 2012; Nagai et al., 2006; Zhao et al., 2014). Recently, it has been shown that repetitive challenges of lipopolysaccharide (LPS), a component of gram-negative bacteria causes HSC dysfunction through direct activation of TLR4 signaling and the resulting proliferative stress (Takizawa et al., 2017).

Humans are estimated to carry approximately 0.2kg of co-existing commensal bacteria amounting to ca. 4x10^13^ cells, which is comparable to the number of cells making up the human body (Sender et al., 2016). The commensal bacteria, often referred to as the microbiota, have been shown to affect many aspects of human health, and its dysregulation leads to many diseases such as obesity, diabetes, atherosclerosis, and inflammatory bowel disease (IBD) (Wang et al., 2017). Recently, accumulating evidence suggests that the microbiota or their components regulates steady-state hematopoiesis through hematopoietic and non-hematopoietic sensing mechanisms, as germ-free conditions or microbiota depletion with antibiotics impairs the development and maintenance of the hematopoietic and immune system, and ultimately increases susceptibility to infection (Balmer et al., 2014; Iwamura et al., 2017; Khosravi et al., 2014; Lee et al., 2019; Yan et al., 2018). However, it remains largely unknown how the hematopoietic system manages substantial microbial infiltration and the associated inflammation under pathophysiologic conditions such as in colitis, and what differences in hematopoietic response exist between acute and chronic settings.

In this study, we employed the colitis mouse model, where administration of dextran sulfate sodium (DSS) has been reported to induce acute and chronic gut inflammation with microbial infiltration, and which resembles the pathogenesis of human IBD (Perse and Cerar, 2012; Wirtz et al., 2017). Here, acute gut inflammation was found to expand HSCs, MPPs and myeloid-committed progenitors in the BM and enhance myelopoiesis via upregulation of granulocyte-macrophage colony stimulating factor (GM-CSF) ligands in local tissues and its receptor on HSPCs. Furthermore, MPPs migrate to the mesenteric lymph node (MLN), a gut-associated lymph node inflamed upon gut inflammation, and differentiate into Ly6C^+^/G^+^ myeloid cells. These processes are mediated by innate immune signals that sense a specific type of microbiota, *Bacteroides*, as antibiotics treatment or genetic ablation of innate immune signaling molecules cancel the DSS-induced hematopoietic responses. The depletion of Ly6C^+^/G^+^ myeloid cells during colitis exacerbates the pathogenesis and implies their role in gut tissue repair. In contrast to acute colitis, chronic gut inflammation hampers HSC function by reducing their repopulation ability upon transplantation. Expansion and migration of HSPCs observed in aged mice with a leaky gut were suppressed in germ-free aged mice. Our findings highlight a cross-organ communication that sense and translates microbial signals to drive myelopoiesis for intestinal tissue repair, whereas chronic microbial insult reduces functionality of long-lived HSCs, which might be relevant to aging-associated chronic disorders.

## Results

### Acute gut inflammation expands HSPCs and skews hematopoiesis towards myelopoiesis

To examine how acute gut inflammation affects early hematopoiesis, wild-type (WT) mice were treated with or without DSS-containing drinking water (DSS or ddW) for 7 days (Fig. 1A). One week of DSS treatment was sufficient to induce acute colitis characterized by shortening of the colon length and the transient loss of body weight (BW), red blood cell (RBC) counts and hemoglobin (Hb) levels in the peripheral blood (PB) due to atrophy, diarrhea and bleeding (Fig. S1A). FACS analysis one day after DSS treatment at day 8 showed a significant increase of HSC^LT^ (long-term HSC defined as CD150^+^CD48^-^Lin^-^Sca-1^+^Kit^+^), MPP2 (multipotent progenitor 2, CD150^+^CD48^+^Lin^-^Sca-1^+^Kit^+^) and MEP (megakaryocyte-erythrocyte progenitor, CD34^-^CD16/32^low^Lin^-^Sca-1^-^Kit^+^) cells in the BM of DSS-treated mice compared to ddW-treated control mice, whereas common lymphoid progenitors (CLPs, Lin^-^c-Kit^low^IL-7R*α*^+^) decreased in DSS-treated BM (Fig. 1B and Fig. S1B). Similarly, an expansion of HSC^LT^ and MPP3/4 (multipotent progenitor 3/4, CD150^-^ CD48^+^Lin^-^Sca-1^+^Kit^+^) but also LSK (Lin^-^Sca-1^+^Kit^+^), LK (Lin^-^Sca-1^-^Kit^+^), MEP, CMP (common myeloid progenitor, CD34^+^CD16/32^low^LK) and GMP (granulocyte-macrophage progenitor, CD34^+^CD16/32^+^LK) cells were observed in the spleen of colitic mice, and HSC^LT^ and MPP2 in the peripheral blood (PB) (Fig. 1B and Fig. S1C-E). Next, the gut-associated mesenteric lymph node (MLN), an inflammatory site underneath the gut tissue was similarly analyzed. Unexpectedly, upon acute colitis, MPP2 and MPP3/4 significantly increased in inflamed MLNs but not in non-inflamed peripheral lymph nodes such as the axillary and inguinal lymph nodes (called PLN thereafter). These results suggest expansion and migration of BM HSPCs preferentially to the inflammatory lymphoid organs (Fig. 1C-D and Fig. S1F). Of note, this phenomena were not specific to chemical-induced mouse colitis, as HSPCs in the MLN were also confirmed in human IBD patients (Fig. S1H-I). Analysis of time-course kinetics revealed a transient expansion of both HSC^LT^ and MPP2 starting at day 5 during DSS treatment and plateauing by day 8 in the BM, while MPP2 in the MLN continued to increase until day 10 (Fig. 1E and Fig. S1G). To determine whether the increase of HSPCs in the BM and MLN resulted from their proliferation, EdU, a DNA analogue was injected 24 hours one day before analysis at day 6 and 7 during ddW/DSS treatment, and its uptake into synthesized DNA strands was measured. The percentage of EdU^+^ cells in each subpopulation of HSPCs was higher in both the BM and MLN of DSS-treated mice than controls at day 7 to 8, suggesting a higher proliferation rate of HSPCs upon acute colitis (Fig. 1F-G). To assess the hematopoietic differentiation capacity of cells found in the MLN, a myeloid colony forming unit (CFU) assay was performed at day 8 (Fig. 1H-I). DSS-treated MLN generated more myeloid colonies than controls, the majority of which were granulocytes. The frequency of MLN-derived CFU cells was ca. 1/1200 that of the BM at steady-state and increased roughly 4-fold after DSS treatment. These results indicate that in response to acute gut inflammation, HSPCs, especially HSC^LT^ and MPP2 expand and skew the hematopoietic program towards myelopoiesis in the BM at the cost of lymphopoiesis, followed by their expansion in the MLN.

**Figure 1.**
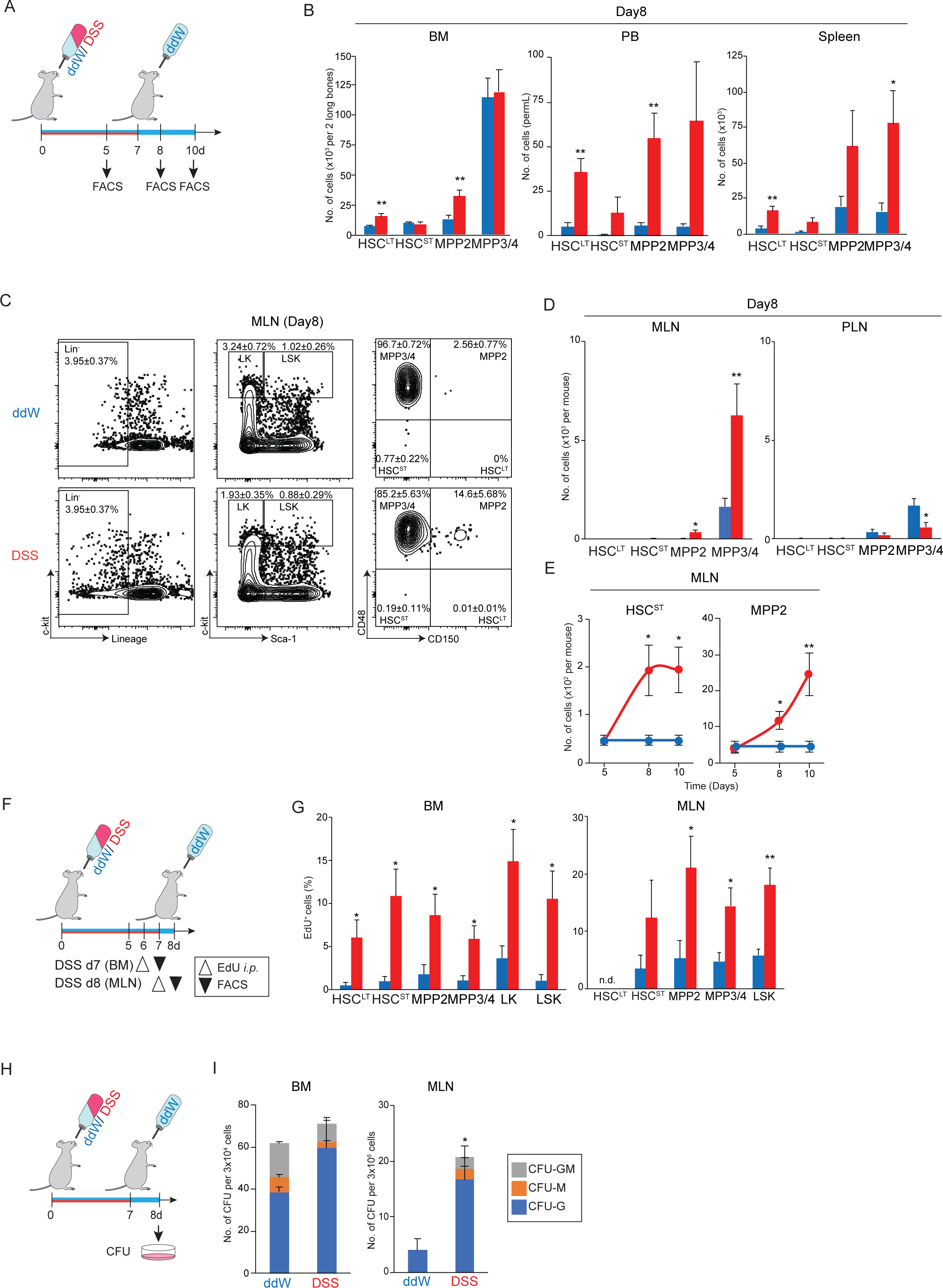
Acute colitis drives myelopoiesis by expanding HSPCs in the BM and MLN. (A) Experimental scheme of DSS-induced acute colitis. (B) Absolute number of HSPCs in the BM, PB and spleen of ddW (blue)- or DSS (red)-treated mice at day (d)8 (1 day after DSS treatment) (n=6-11 from 2-4 experiments). (C) Representative FACS plots of HSPCs in the MLN from ddW (upper)- or DSS (lower)- treated mice at d8. (D) Absolute number of HSPCs in the MLN and PLN of ddW (blue)- or DSS (red)-treated mice (n=5-9 from 2-4 experiments). (E) Time-course analysis of HSPC number in the MLN at d5, d8, and d10 (n=3-5 from 2 experiments). (F) Experimental scheme of EdU assay: WT mice treated with ddW (blue) or DSS (red in G) for 7 days were *i.p.* injected with 100μg of EdU at d6 or d7 and analyzed the following day. (G) Percentage of EdU^+^ HSPCs in the BM and MLN (n=4 from 2 experiments). (H) Experimental scheme for colony formation assay. BM and MLN cells isolated at d8 were cultured *in vitro* for 14 days until analysis. (I) Number of colonies and colony types derived from the BM (left) and MLN (right) (n=3-6 from 3 experiments). n.d., not detected; \**p*<0.05; \*\**p*<0.01 (two-tailed t-test).

### Upregulation of GM-CSF and its ligand regulates MPP2 expansion upon acute gut inflammation

To understand the molecular mechanism underlying gut inflammation-induced HSPC activation, RNA sequencing was performed on MPP2 sorted from the BM and MLN one day post ddW/DSS treatment at day 8 (Fig. 2A). The sequencing data shows drastic transcriptional changes between MPP2 of the BM and MLN (Fig. 2B-C), where the expression of 80 genes in the BM and 215 genes in the MLN changed significantly and differentially, many of which are known to be involved in inflammation, cell differentiation, migration, and cellular metabolism. Amongst the genes, *Csf2rb2* and *Myl10* commonly changed in MPP2 of both DSS-treated BM and MLN, while *Csf2rb2* and *Csf2rb*, both coding for GM-CSFR (granulocyte-macrophage colony-stimulating factor receptor) were most significantly upregulated in the BM (Fig. 2D). The upregulated expression of GM-CSFR and was confirmed at the transcriptional and protein level with quantitative PCR and FACS analysis (Fig. 2E-F). ELISA showed upregulation of GM-CSF protein also in the BM fluid and MLN lysate upon DSS treatment, but not in the plasma (Fig. 2G). In agreement with the previous study (Griseri et al., 2012), these data suggest that acute gut inflammation induces upregulation of GM-CSFR expression on HSPCs and its ligand locally produced in the BM and MLN, which might cause the expansion of HSPCs.

**Figure 2.**
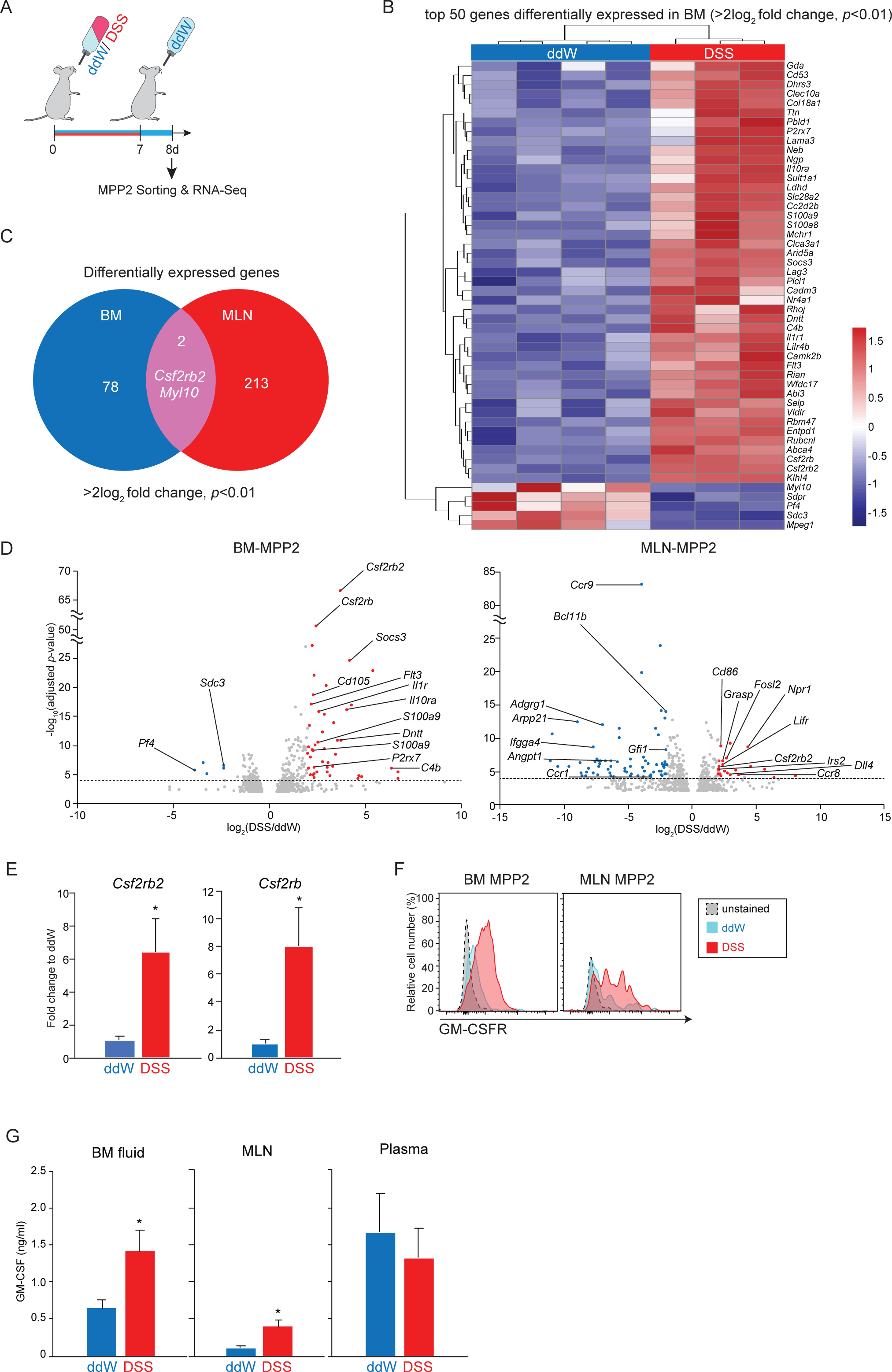
Acute colitis activates GM-CSF signals in HSPCs for their expansion. (A) Experimental scheme of RNA-seq with 100 MPP2 (LSKCD48^+^CD150^+^) cells FACS-sorted from the BM or MLN of ddW/DSS-treated WT mice at d8. (B) A heatmap of top 50 DEGs of MPP2 from DSS-treated BM compared to ddW-treated BM (n=3-4, >2log_2_ fold change, *p*<0.01). The color code represents expression levels of each gene as indicated. (C) Venn diagram of DEGs between MPP2 in the BM (blue) and MLN (red) (>2log_2_ fold change, *p*<0.01). (D) Volcano plots of DEGs of MPP2 in the BM (left) and MLN (right). Red and blue dots represent upregulated and downregulated genes, respectively (>2log_2_ fold change, *p*<0.0001). (E) qPCR analysis of *Csf2rb2* and *Csf2rb* genes in MPP2 from ddW (blue) and DSS (red) BM (n=3 from 2 experiments). (F) Representative FACS histogram of GM-CSFR expression by MPP2 cells in the BM and MLN (DSS, red; ddW, blue; unstained, gray). (G) ELISA of protein levels of GM-CSF in the BM fluid, MLN lysate, and plasma of mice treated with ddW (blue) and DSS (red) (n=3-4 from 2 experiments). \**p*<0.05 (two-tailed t-test).

### Acute gut inflammation-induced HSPC expansion is mediated by innate immune signaling

We previously reported that LPS, a cellular component of gram-negative bacteria directly activates HSCs via TLR-4 (Takizawa et al., 2017). Given that the microbiota, 9-47% of which consists of gram-negative bacteria spreads throughout the body upon gut inflammation (Lagier et al., 2012), we hypothesized that they would be sensed by innate immune recognition receptors such as TLRs. Therefore, *Trif*^-/-^;*Myd88^-/-^* -[double knock out mice (DKO), which lacks TLR and IL-1R-mediated innate immune signaling were next analyzed (Akira and Takeda, 2004). While DSS treatment expanded HSPCs (HSC^LT^/MPP2) in the BM of WT mice (Fig. 1B), the effect was completely abrogated in DKO mice (Fig. 3A-C). Similarly, MPP expansion in the MLN and spleen was seen only in WT and not in DKO mice (Fig. 3D-E and Fig. S2A). In contrast, CLPs in the BM decreased in both DSS-treated WT and DKO mice and implies a TLR/IL-1R-independent mechanism as previously reported (Terashima et al., 2016). Moreover, as a consequence of expansion of MPP and myeloid progenitors, we confirmed expansion of Ly6C/G^high/low^ cells in the BM and spleen of DSS-treated WT but not DKO mice (Fig. S2B-D). These results indicate that the observed gut inflammation-induced expansion of HSPCs (HSC^LT^/MPP2) in the BM and MLN is regulated by innate immune signaling.

**Figure 3.**
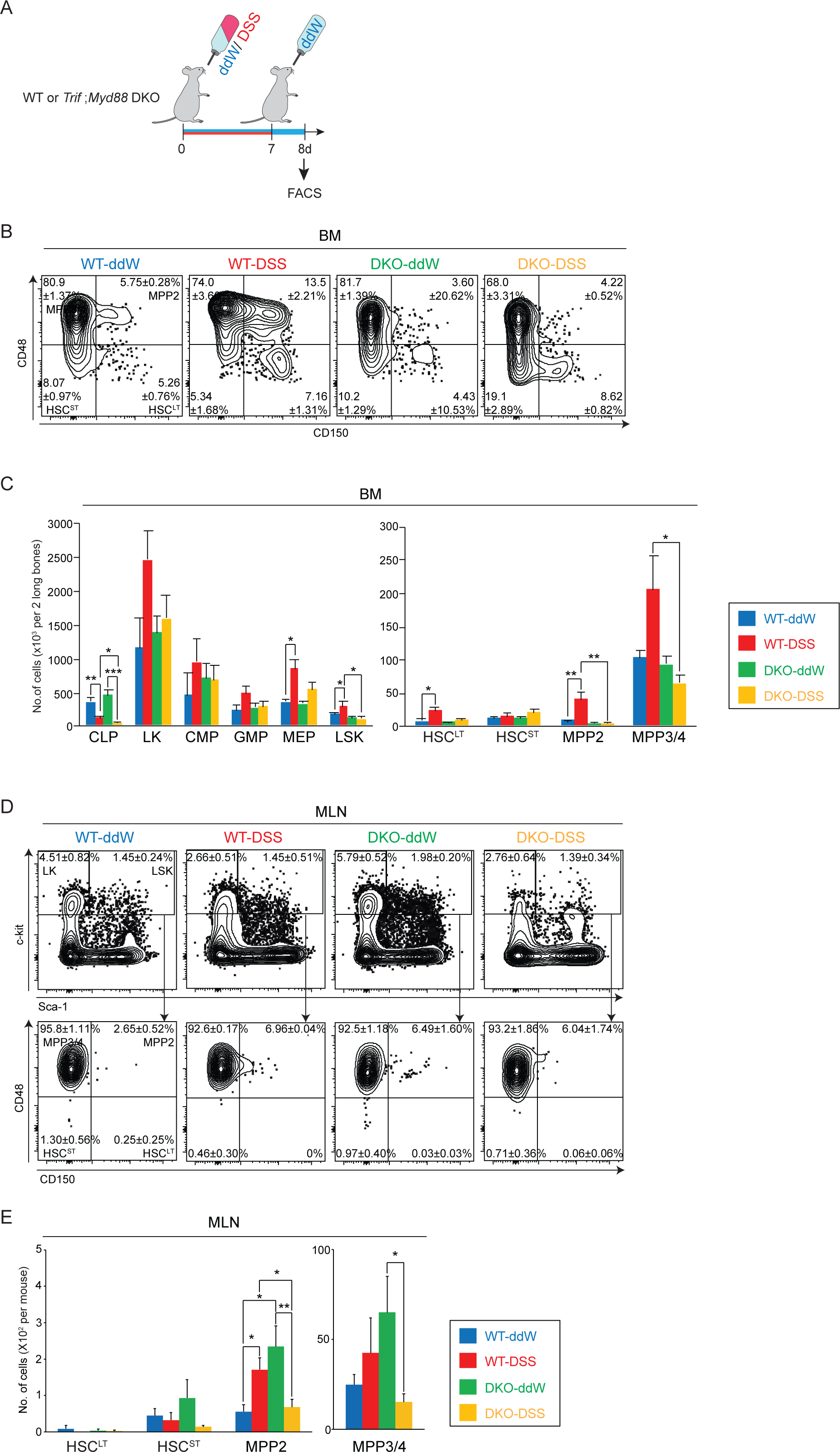
Acute colitis-induced HSPC expansion and migration occurs through innate immune signaling. (A) Experimental scheme of colitis induction in WT or *Trif*;*Myd88* DKO mice. (B) Representative FACS plots of the BM of ddW- or DSS-treated WT or *Trif*;*Myd88* DKO mice. (C) Absolute number of HSPCs in the BM of WT-ddW (blue), WT-DSS (red), DKO-ddW (green), and DKO-DSS (orange) (n=6-7 from 3 experiments). (D) Representative FACS plots of HSPCs in the MLN of ddW- or DSS-treated WT or *Trif*;*Myd88* DKO mice. (E) Absolute number of HSPCs in the MLN of WT-ddW (blue), WT-DSS (red), DKO-ddW (green), and DKO-DSS (orange) (n=4-7 from 3 experiments). \**p*<0.05; \*\**p*<0.01; \*\*\**p*<0.001 (two-tailed t-test).

### Gut-derived *Bacteroides* promotes HSPC expansion and differentiation into myeloid cells

Assuming that the innate immune signaling activated by acute colitis is mediated by infiltrating microbiota, we next sought to determine the species of gut microbiota responsible for the DSS-induced changes in hematopoiesis. WT mice were treated with the single antibiotics, neomycin (NM), metronidazole (MNZ), vancomycin (VCM), or ampicillin (ABPC), each acting against a distinct spectrum of microorganisms. Following 4 weeks of antibiotics pretreatment, mice were challenged with ddW/DSS for one week and analyzed on day 7 (Fig. 4A). NM- and MNZ-pretreated mice showed DSS-induced MPP expansion similar to non-pretreated mice (nTx) in the BM and to a greater degree in the MLN, while MPP expansion was completely suppressed by pre-treatment with VCM and ABPC (Fig. 4B). Results suggest that HSPC expansion in the BM and MLN is regulated by a specific type of microbiota unresponsive to NM and MNZ but sensitive to VCM and ABPC. We next isolated gut contents from antibiotics-pretreated mice and extracted DNA to perform metagenome analysis. When mice treated with or without DSS were compared, unweighted and weighted UniFrac analysis showed that DSS treatment did not affect microbiome composition, while pretreatment with each antibiotics resulted in significant changes to the gut microbial composition as expected (Fig. 4C and Fig. S3A). Especially, ABPC-pretreated mice with fewer color combinations, indicating decreased microbiota diversity showed higher mortality post DSS treatment compared to nTx or mice pre-treated with the other antibiotics (Fig. S3B). This finding may imply the role of microbiota diversity in tissue protection or anti-inflammation responses against IBD. We next screened for HSPC-regulating candidate bacteria that increased in NM- and MNZ-pretreated mice and decreased in VCM- and ABPC-pretreated mice, and found that s24-7 and Bacteroidaceae, which are both a family of gram-negative Bacteroidales bacteria, showed a positive correlation with HSPC number as shown in Fig. 4B (Fig. 4C). To validate whether Bacteroidales induces HSPC expansion, cell lysate derived from 6 species of Bacteroidales (*Bactroides*) was *i.p.* injected into WT mice without DSS treatment and analyzed 12 hours later (Fig. 4D). Systemic injection of the lysate induced HSPC expansion in the BM and MLN that recapitulated the DSS-induced effect (Fig. 4E and Fig. S3C). However, when the lysate was injected into *Trif*^-/-^;*Myd88^-/-^* mice or upon injection of *Faecalibacterium*, another family of gram-negative bacterium, no MPP response was observed. These results demonstrate that the colitis-induced expansion of HSPCs is regulated by *Bacteroides* that activates innate immune signaling.

**Figure 4.**
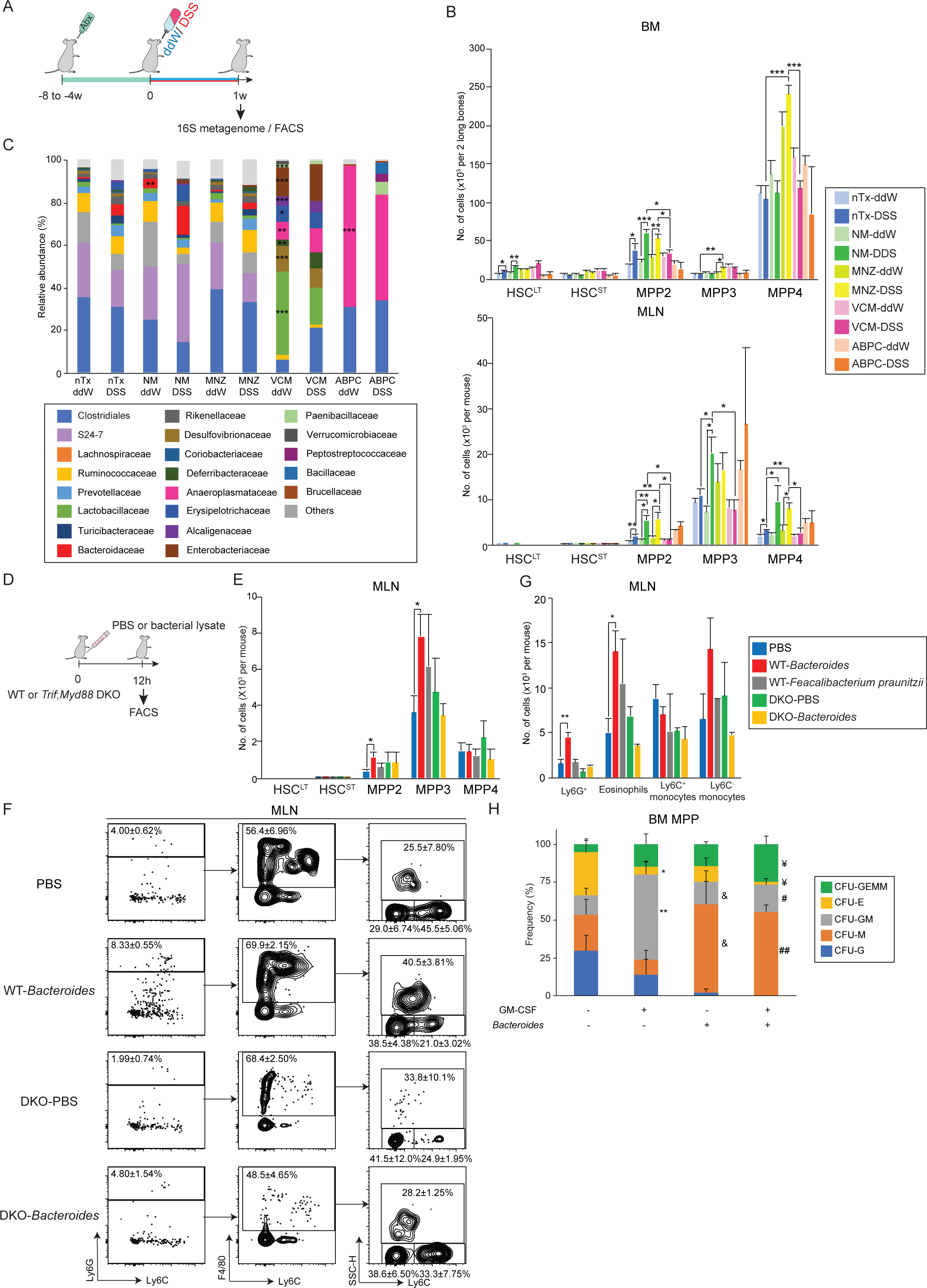
*Bacteroides* regulates HSPC expansion and migration through innate immune sensing in response to acute colitis. (A) Experimental scheme of antibiotics pretreatment followed by colitis induction: 4 week-old WT mice were treated with or without the following antibiotics, nTx (non-treated), NM, MNZ, VCM, and ABPC for 4-8 weeks, followed by ddW/DSS and immediate FACS and metagenome analysis. (B) Absolute number of HSPCs in the BM (top) and MLN (bottom) of nTx-ddW/DSS (light/dark blue), NM-ddW/DSS (light/dark green), MNZ-ddW/DSS (dark/light yellow), VCM-ddW/DSS (light/dark pink), and ABPC-ddW/DSS (light/dark orange) mice (n=2-12 each from 2 experiments). (C) Relative abundance of gut microbiota in mice pretreated with or without antibiotics. Each color represents a specific family of bacteria as stated in the legend below (n=2-10 for each, from 2 experiments). \**p*<0.05; \*\**p*<0.01; \*\*\**p*<0.001 (two-tailed t-test, compared to nTx-ddW). (D) Experimental scheme of bacterial cell lysate injection: WT or *Trif*;*Myd88* DKO mice were *i.p.* injected with PBS, *Bacteroides* or *Feacallibacterium praunitzii* lysate (15μg), and MLN cells were analyzed 12 hours later. (E) Absolute number of HSPCs in the MLN of WT or DKO mice injected with PBS (blue/green), *Bacteroides* (red/yellow) or *Feacalibacterium praunitzii* (gray) (n=3-4 from 3 experiments). \**p*<0.05 (F) Representative FACS plots and (G) absolute number of myeloid cells in the MLN of WT or *Trif*;*Myd88* DKO mice injected with PBS (blue/green), *Bacteroides* (red/yellow) or *Feacalibacterium praunitzii* (gray) (n=3-4 from 3 experiments). *p*<0.05; \*\**p*<0.01. (H) Frequency of colony types 14 days after culture of BM-MPP with or without GM-CSF/*Bacteroides* (n=48 from 3-4 experiments) (GEMM, granulocyte, erythroid, macrophage, and megakaryocyte; E, erythroid; GM, granulocyte-macrophage; M, macrophage; G, granulocyte); *,^$,¥,&,#^*p*<0.05; **,^$$,¥¥,##^*p*<0.01, n.s., not significant (two-tailed t-test, *control vs GM-CSF alone; ^$^control vs *Bacteroides* alone; ^¥^control vs *Bacteroides* and GM-CSF; *^&^*GM-CSF alone vs *Bacteroides* alone; *^#^*Bacteroides alone vs *Bacteroides* and GM-CSF).

To understand the biological consequence of MPP migration to inflammed MLNs, the mature hematopoietic compartment was analyzed after *Bacteroides* lysate injection into WT mice. CD11b^+^Ly6G^+^ cells in the BM decreased as observed in DSS-treated mice, while Ly6G^+^ cells increased in the PB, spleen and together with CD11b^+^Ly6G^-^F4/80^+^SSC^hi^ eosinophils in the MLN (Fig. 4F-G and Fig. S3D-F). In contrast, the effect could not be confirmed in *Bacteroides*-injected *Trif*^-/-^;*Myd88^-/-^* mice. When MPPs from WT BM were *in vitro* cultured with *Bacteroides*, GM-CSFR expression was significantly upregulated two days after culture (Fig. S3G-H). Furthermore, single-cell CFU assay showed *Bactreroides* stimulation shifted MPP differentiation towards macrophage colonies, with or without the addition of GM-CSF while not affecting colony efficiency (Fig. 4H and S3I-L). These results suggest that *Bacteroides* can directly activate MPPs to upregulate GM-CSFR and bias their differentiation towards myeloid cells.

### Blocking HSPC migration to the MLN exacerbated DSS-induced acute gut inflammation

We next examined HSPC localization in the MLN using the *Hlf^tdTomato+^* knock-in reporter mouse that enables the tracking of HSPCs by tdTomato expression inserted under the HSPC-specific *Hlf* promoter (Yokomizo et al., 2019). MLNs imaged at day 7 (Fig. 5A-B) were consistent with FACS data (Fig. 1D), as DSS treatment increased the number of *Hlf^+^* HSPCs in the MLN, the majority of which appeared to be in close proximity to CD31^+^ endothelial cells rather than PNAd^+^ high endothelial venules (Fig. 5B). The distance of *Hlf^+^* cells to the nearest vessel, when quantified showed that more than half (>53%) were localized within 20μm compared to artificially generated random spots (Fig. 5C). Furthermore, time-lapse imaging of *Hlf^+^* HSPCs showed active movement in the vessels upon DSS treatment compared to ddW-treated controls (data not shown). These results suggest that *Hlf^+^* HSPCs may migrate to the MLN through the vasculature upon acute gut inflammation.

**Figure 5.**
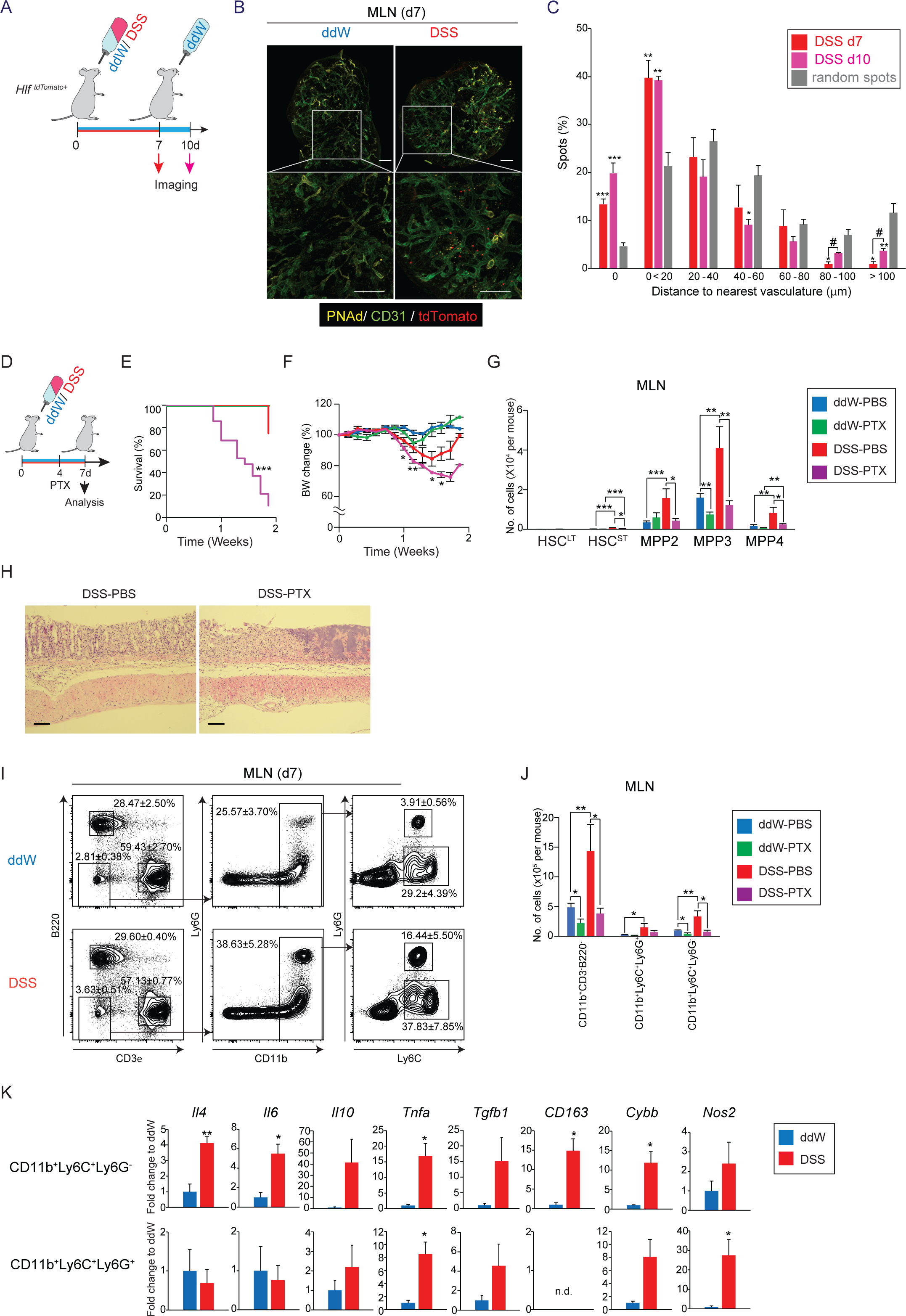
HSPCs migrate to the MLN through vessels and differentiate into anti-inflammatory Ly6C^+^/G^+^ cells upon acute colitis. (A) Experimental scheme: MLN of DSS-treated *Hlf^tdTomato+^* mice was harvested at d7 and d10 for whole-tissue imaging. (B) Representative images of MLN from mice treated with ddW (left) or DSS (right) at d10 (CD31, green; PNAd, yellow; tdTomato, red). Area enclosed by white squares are enlarged in the lower panels (scale bar, 200μm). (C) Tissue distribution of *Hlf^tdTomato+^* HSPCs in relation to CD31^+^ endothelial cells at d7 (red) and d10 (pink). Number of *Hlf^tdTomato+^* HSPCs in the MLN within a certain distance to the nearest vasculature or computationally generated random spots was determined and the percentage of each fraction calculated. (n=51-439 cells per organ from 3-6 experiments). \**p*<0.05, \*\**p*<0.01, \*\*\**p*<0.001, compared to random spots; ^#^*p*<0.05, d7 compared to d10 (two-tailed t-test). (D) Experimental scheme of pertussis toxin (PTX) treatment: WT mice were *i.p.* injected with 1μg of PTX at d4 of DSS treatment. (E) Survival rate post PTX treatment (n=10-40 from 6 experiments). *p<0.05 with Kaplan-Meier estimate. (F) Time-course kinetics of body weight (BW) change post PTX treatment (n=11-39 from 6 experiments). (G) Absolute number of HSPCs in the MLN at d7 (n=4-11 from 2-5 experiments). (H) HE staining of gut tissue sections of DSS-PBS versus DSS-PTX mice. (I) Representative FACS plots and (J) absolute number of mature cell populations in the MLN at d7 (n=4-11 from 2-5 experiments). (K) Expression of anti-inflammatory, MDSC-, and M2 macrophage-related genes in CD11b^+^Ly6C^+^ and Ly6G^+^ cells from ddW- (blue) and DSS- (red) treated MLN (n=3 from 2 experiments). \**p*<0.05; \*\**p*<0.01; \*\*\**p*<0.001.

To understand the role of MPP migration to the MLN in colitis pathogenesis, we attempted blocking their mobilization with pertussis toxin (PTX), which inhibits the *α* subunit of small G proteins. Mice injected with PTX at day 4 during DSS treatment showed high mortality and significant BW loss with slower recovery compared to PBS-injected controls (Fig. 5D-F). The colitis-induced HSPC migration and expansion in the MLN was completely cancelled in PTX-injected mice (Fig. 5G), while gut histology revealed more damage (Fig. 5H). Cell populations that increased upon DSS treatment, but were blocked by PTX were CD11b^+^Ly6C^+^Ly6G^+^ (Ly6C^+^/G^+^) and CD11b^+^Ly6C^+^Ly6G^-^ (Ly6C^+^/G^-^) myeloid cells (Fig. 5I-J). These results suggest that Ly6C^+^/G^+^ and Ly6C^+^/G^-^ myeloid cells might suppress gut tissue damage. We next checked for anti-inflammatory cells such as myeloid-derived suppressor cells (MDSCs), a population characterized by the surface antigens CD11b and Ly6C/G that arises in the case of cancer and other diseases with immunosuppressive activity (Gabrilovich and Nagaraj, 2009). Gene expression analysis of Ly6C^+^/G^-^ cells of DSS-treated MLN revealed upregulation of *Il6*, which is required for the expansion and activation of myeloid-derived suppressor cells (MDSCs) (Marigo et al., 2010) and their suppressive activity (Chalmin et al., 2010). The TH2-type cytokines, *Il4 and Il10*, both required for the resolution of an immune response (Poe et al., 2013), and the latter recently reported to ameliorate colitis through its secretion by MDSCs (Zheng et al., 2021) were also upregulated. *Tnfa* (Condamine et al., 2015), *Cybb* (Corzo et al., 2009), and *Nos2* (Raber et al., 2012), all of which are associated with the immunosuppressive action of MDSCs via reactive oxygen species and nitric oxide secretion (Veglia et al., 2018)(Fig. 5K) and the anti-inflammatory M2 macrophage-related gene, *Cd163* highly expressed during the resolution of inflammation (Yang et al., 2016) were among the others significantly upregulated in Ly6C^+^/G^+^ and Ly6C^+^/G^-^ cells. Of note, expansion of their human counterparts, myeloid cells such as CD33^+^, CD14^+^ and CD16^+^ cells were also confirmed in the MLN of human IBD patients (Fig. S1J-K). These results suggest that HSPCs migrating through mesenteric vessels generate immunosuppressive Ly6C^+^/G^+^ cells in the MLN that may cooperate in tissue repair against DSS-induced colitis.

### Ly6C^+^/G^+^ cells generated locally in MLN contribute to intestinal tissue repair

We next assessed the functional role of Ly6C^+^/G^+^ cells in the pathogenesis of colitis. To determine the optimal window for cell depletion, time-course kinetics of the mature compartment was analyzed. Unlike the BM, the MLN showed an increase of eosinophils and Ly6C^+^ monocytes at day 8 which was sustained thereafter, while the increase of Ly6G^+^ cells and Ly6C^-^ monocytes was transient (Fig. 6B-C). Although the BM showed increased Ly6C^+^ cells and decreased Ly6G^+^ cells, PB analysis showed decreased eosinophils and Ly6C^-^ cells with constant Ly6G^+^ cells over time (Fig. S4A-C), and implies the local generation of Ly6C^+^/G^+^ cells in the MLN. Mice were injected with a neutralizing antibody against the Gr-1 antigen to deplete both Ly6G^+^ and Ly6C^+^ cells (Carr et al., 2011), at day 6 and day 7 during DSS treatment (Fig. 6D). The efficiency of cell depletion reached almost 100% for Ly6G^+^ cells, and 50% for Ly6C^+^ cells in the BM and MLN, respectively (Fig. S4F). Under these conditions, injection of the Gr-1 antibody into DSS-treated mice significantly shortened colon length, which is a sign of tissue atrophy compared to mice treated with DSS plus an isotype-matched antibody (IgG) (Fig. 6E). Consistently, several pathological parameters such as BW loss and the disease activity index (DAI) showed exacerbation in the Gr-1 antibody-injected group (Fig. 6F and Fig. S4D). In addition, histology confirmed that mice treated with DSS and the Gr-1 antibody developed severe inflammation with complete loss of Goblet cells, infiltration of lymphocytes and the destruction of epithelial surfaces, resulting in an overall higher histological score in contrast to isotype-matched controls (Fig. 6G-H). Immunophenotyping of early hematopoietic cells following antibody injection revealed that Ly6C^+^/G^+^ cell depletion results in greater expansion/migration of MPP2 to the MLN which was not observed in the BM (Fig. S4E). This suggests that MPP may sense the local demands for Ly6C^+^/G^+^ cell generation and differentiate in the MLN following acute colitis and cell depletion. Collectively, these data indicate that MPPs in the MLN locally differentiate into Ly6C^+^/G^+^ cells to promote the repair of gut tissue upon colitis.

**Figure 6.**
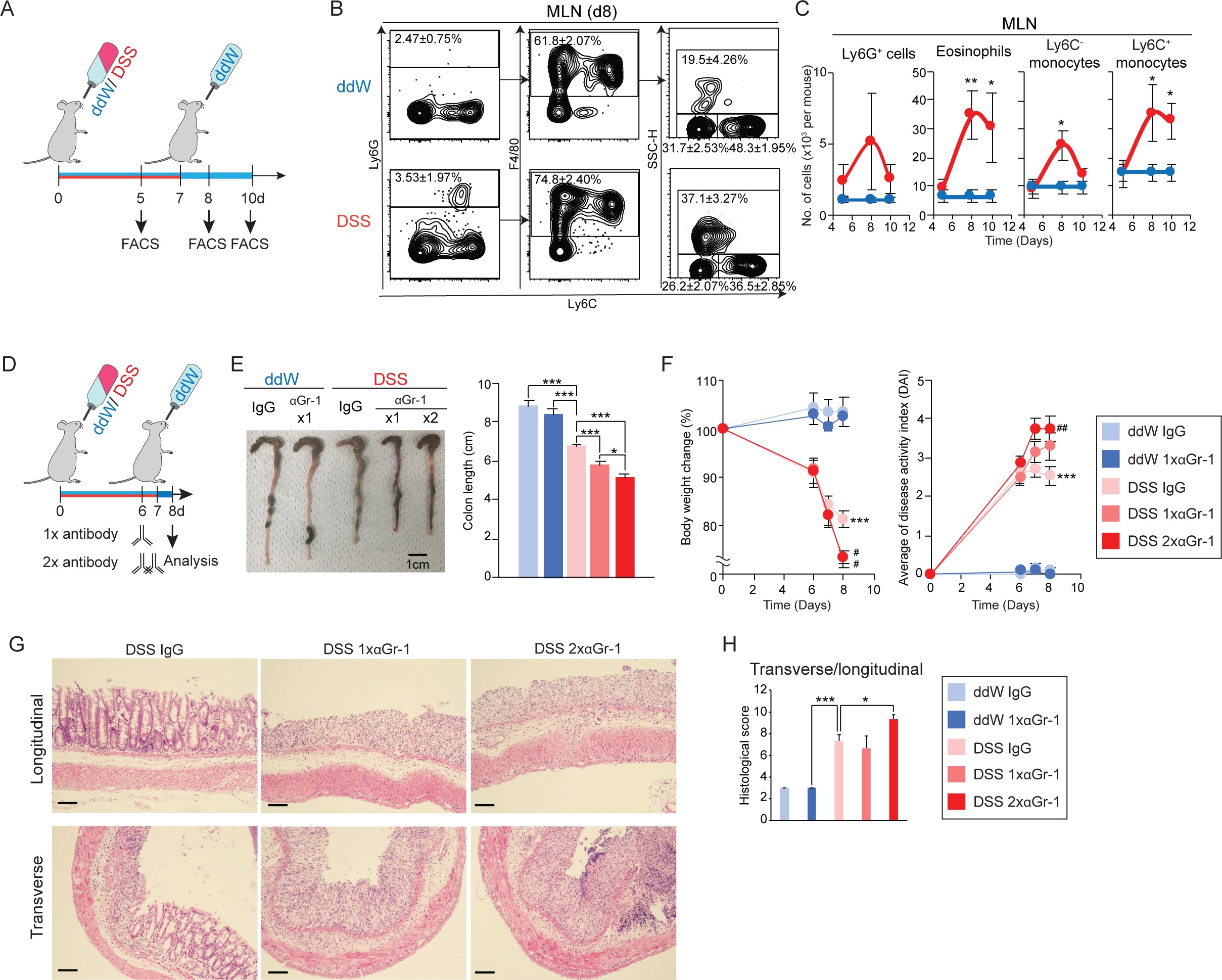
Acute colitis expanded Ly6C^+^/G^+^ cells in MLN contributes to gut tissue repair. (A) Experimental scheme of time-course FACS analysis at d5, d8, and d10. (B) Representative FACS plots and (C) time-course kinetics of myeloid cell populations in d8 MLN pre-gated on B220^-^CD3*ε*^-^CD11c^-^ cells. (n=3-5 from 2 experiments). (D) Experimental scheme of Ly6C^+^/G^+^ cell depletion: WT mice were *i.p.* injected with 500μg of neutralizing anti-Gr-1 antibody (*α*Gr-1) or IgG isotype-matched control (IgG) at d6 (single dose) or d6 and d7 (double dose). (E) Representative images of the colon (left) and measurements of colon length (right). (F) Time-course analysis of BW loss (left) and disease activity index (DAI) (right) were monitored daily for up to 8 days (ddW IgG, light blue; ddW *α*Gr-1, dark blue; DSS IgG, light pink; DSS 1x*α*Gr-1, dark pink; DSS 2x*α*Gr-1, red) (n=4-7 from 3 experiments). (G) HE staining of longitudinal (upper) and transverse (lower) gut tissue sections and (H) histological scoring of transverse and longitudinal sections at d8. Scale bars represent 100µm. \*\*\**p*<0.001; ^#^p<0.05; ^##^p<0.01 (two-tailed t-test, *ddW-IgG vs DSS IgG; ^#^DSS IgG vs DSS 1x*α*Gr-1 or DSS 2x*α*Gr-1). (J) n=3, \**p*<0.05; \*\**p*<0.01; \*\*\**p*<0.001.

### Chronic gut inflammation impairs HSC self-renewal and immune-reactiveness

Finally, we examined the effects of chronic gut inflammation on early hematopoiesis to explore any differences in hematopoietic response compared to acute conditions. A cycle of one week ddW/DSS treatment and one week ddW treatment was repeated 3 times to induce chronic colitis as previously reported (Fig. 7A) (Wirtz et al., 2017). Analysis of PB parameters and BW changes showed decreased RBC counts, Hb and BW following chronic colitis, while white blood cell (WBC) and platelet (PLT) counts increased with all parameters recovering to normal levels by 3 months post colitis induction (Fig. S5A). To first test for HSC functionality, BM cells from chronic ddW/DSS-treated mice were serially transplanted into WT recipients followed with a monthly check of PB donor chimerism and terminal BM analysis (Fig. 7A). Results showed significantly lower multilineage hematopoietic reconstitution in the PB of transplants with chronic DSS-treated BM cells, while BM donor chimerism of HSCs, MPPs and mature cells were all reduced compared to controls (Fig. 7B-C and Fig. S5B). Secondary recipients showed significantly decreased myeloid cells in the PB and HSPCs in the BM, and lower survival rates presumably due to BM failure (Fig. S5C-F). Next, the immune response of chronically-inflamed HSCs was assessed via an LPS re-challenge experiment (Fig. S5G) as LPS has been previously reported to induce HSC mobilization (Cline and Golde, 1977; Takizawa et al., 2017). BM analysis 16 hours after LPS injection showed decreased HSC^LT^ numbers in ddW-treated control mice due to mobilization, while chronic DSS-treated mice showed little reduction (Fig. S5H). These results demonstrate that chronic gut inflammation impairs hematopoietic repopulating ability and the immune response of HSCs in a cell-intrinsic manner.

**Figure 7.**
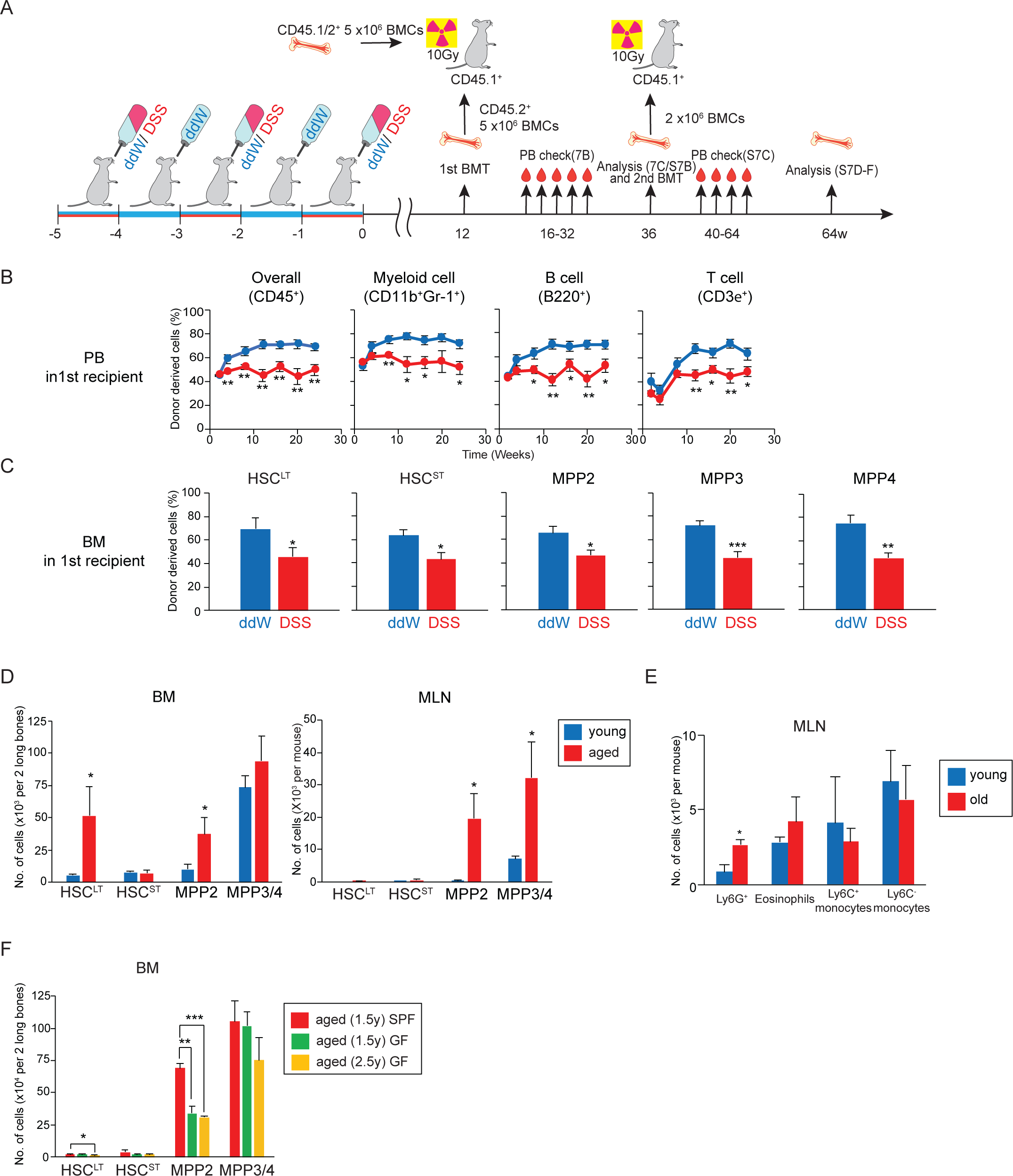
Chronic colitis impairs HSC transplantability and immune-reactivity. (A) Experimental scheme of chronic colitis and serial BM transplantation (BMT): WT mice (CD45.2^+^) were treated with one week of ddW (blue)/DSS (red) and one week of ddW for 3 cycles. At 3 months, 5x10^6^ total BM cells were co-transplanted together with 5x10^6^ total BM competitor cells. Donor chimerism in the PB was assessed monthly for up to 20 weeks. (B) At 24 weeks post 1^st^ BMT, total BM cells isolated from 1^st^ recipients were analyzed (C) and serially transplanted into lethally irradiated WT mice (CD45.1^+^), followed by donor chimerism check (Fig. S7). (B) Time-course kinetics of donor chimerism in overall (CD45^+^), myeloid (CD11b^+^Gr-1^+^), B (B220^+^), and T (CD3*ε*^+^) cells in the PB of 1^st^ recipients (n=8-12 from 2 experiments). (C) Donor chimerism in HSPCs in the BM of 1^st^ recipients (n=8-12 from 2 experiments). (D) Absolute number of HSPCs and (E) myeloid cells in the BM (left) and MLN (right) of young (8-16 weeks-old, blue) and aged (>100 weeks-old, red) WT mice (n=3-4 from 3 experiments). (F) Absolute number of HSPCs in the BM of aged mice under specific-pathogen-free (SPF) conditions (1.5 years-old, red) or germ-free (GF) conditions (1.5 year-old, green; 2.5 year-old, yellow) (n=3 from 2 experiments). \**p*<0.05;\*\**p*<0.01; \*\*\**p*<0.001 (two-tailed t-test).

Finally, given that aging causes a leaky gut and thus readily allows microbial translocation through the epithelial barrier (Jeong et al., 2017), we tested whether the hematopoietic changes observed upon acute colitis is specific to DSS-induced gut inflammation or a general mechanism in other inflammatory contexts such as inflammaging (Kovtonyuk et al., 2016). FACS analysis of aged and young WT mice showed expansion of MPP and Ly6G^+^ cells only in aged BM and MLN (Fig. 7F-G). Furthermore, MPP expansion observed in specific-pathogen-free (SPF) mice significantly diminished in the BM of germ-free (GF) mice, suggesting that MPP expansion and Ly6G^+^ cell differentiation is mediated by the microbiota (Fig. 7H).

## DISCUSSION

Here we report that upon acute gut injury, *Bacteroides* infiltrates the body to stimulate innate immune receptors such as TLRs and induces the expansion of HSPCs in the BM via GM-CSF signals. Subsequently, MPPs migrate to inflamed MLNs through vessels where they will further differentiate into Ly6C^+^/G^+^ myeloid cells that contribute to gut tissue repair. In contrast, chronic gut inflammation dramatically impairs HSC function by reducing their reconstitution potential and weakening immune responses to subsequent immunological challenges. These findings highlight the importance of tissue surveillance by the hematopoietic system equipped with a sensing mechanism to recognize peripheral tissue-derived damage signals and to integrate them into the hematopoietic program to deal with microbial infiltration and tissue damage at local sites. However, excessive inflammation in turn damages hematopoietic function, supporting the notion that acute inflammation brings about beneficial effects whereas chronic is more detrimental.

The DSS-induced colitis model used in this study recapitulates characteristics of the human IBD disease (Wirtz et al., 2017). Our preliminary data suggests that human IBD patients likewise show localization of early hematopoietic cells at the MLN such as MPPs (Lin^-^hCD34^+^hCD38^-^hCD45RA^-^hCD90^-^) and multi-lymphoid progenitors (MLPs) (Lin^-^hCD34^+^hCD38^-^hCD45RA^-^hCD90^-^) (Fig. S1H-I), and can probably be reproduced by the same type of microbial species as *Bacteroides* is one of the key microbiota with negative correlation to the clinical score of Crohn’s disease (Khosravi et al., 2014). *Faecalibacterium*, another gram-negative bacteria had no impact on HSPC and myeloid cell expansion (Figure 4D-G), suggesting that not all microbiota associated with IBD are able to boost hematopoietic responses to promote tissue repair. This is possibly because microbial components like LPS are species-specific, and elicit different biological reactions in the host. Indeed, the molecular structure of *Bacteroides*-derived LPS is distinct from that of *Escherichia coli* and fails to incite the production of inflammatory cytokines, e.g., interleukin (IL)-10, tumor necrosis factor alpha (TNF*α*) and immune tolerance in innate immune cells (Watson and Kim, 1963). Moreover, the induction of T1D seems to be mediated by the microbiota that stimulate TLR-MyD88-dependent pathways (Wen et al., 2008), whereas TRIF-dependent pathways provide protective signals against T1D (Burrows et al., 2015).

Several groups have recently characterized changes in the microbiome as key regulators of normal and diseased hematopoiesis. The imbalance of gut microbiota caused by antibiotics or germ-free conditions result in hematopoietic dysregulation at different levels (Yan et al., 2018). One study showed that antibiotics treatment diminishes HSPCs in the BM through signal transducer and activator of transcription (STAT) 1, but not the MyD88- and nucleotide-binding oligomerization domain-containing protein (NOD) 2-dependent pathway (Josefsdottir et al., 2017). The microbiota, sensed by NOD1 expressing MSCs or MyD88 expressing CX3CR1^+^ mononuclear cells in the BM regulate HSPC pool size through proinflammatory cytokines (Iwamura et al., 2017; Lee et al., 2019) but can also cause the malignant transformation of HSCs as high fat diet-induced anemia and myeloproliferative neoplasm-like disease in *Spred1^-/-^* mice was suppressed by antibiotics-mediated microbiota depletion (Tadokoro et al., 2018). Our study demonstrates that under pathological conditions like colitis, the detection of infiltrating *Bacteroides* is dependent on MyD88- and TRIF-mediated innate immune signals (Fig. 4E). Although MyD88 and TRIF transduce signals through TLR and IL-1R, based on our in vitro data, TLR but not IL-1R is likely to detect *Bacteroides* and regulate HSPC behavior (Fig. 4H), as IL-1*β* ligand did not appear upregulated in the BM fluid, MLN nor plasma (data not shown). Whether *Bacteroides* directly stimulates HSPCs through TLR or indirectly via cytokines expressed by non-hematopoietic cells (e.g., MSCs or *Bacteroides*-derived molecules such as proteins and metabolites), and acts as a key driver for early hematopoietic regulation remains to be determined.

Several cytokines (e.g., IFN, IL-1, TNF, IL-6, GM-CSF, etc.) are known to expand HSPCs in inflammatory diseases. Here, gut inflammation locally upregulated GM-CSF and its receptor in the BM to regulate HSPC expansion (Fig. 2E-G). This is in agreement with a previous finding that reports the reduction of myeloid progenitors by blocking GM-CSF (Griseri et al., 2012), although compared to our results, this study found that the GM-CSF-induced accumulation of myeloid progenitors in the spleen and colon exacerbates colitis, indicative of their pathologic role in disease progression. On the contrary, others have shown that genetic ablation of GM-CSF increases susceptibility to DSS-induced acute colitis and that GM-CSF administration improves recovery from colitis, suggesting the protective role of GM-CSF (Sainathan et al., 2008; Xu et al., 2008). It is possible that GM-CSF regulates not only HSPC expansion but also their migration to the MLN as has been previously shown for leukocytes (Khajah et al., 2011). These contradictory results regarding the effect of GM-CSF may stem from the different experimental models tested. Griseri and colleagues the acquired immunity-driven colitis model involving the adaptive transfer of naïve T cells (CD4^+^CD25^-^CD45RB^hi^), whereas the DSS-induced colitis model predominantly relies on innate immunity (Wirtz and Neurath, 2007). It may well be that the same cytokine can exert different outcomes on the pathogenesis, given that the underlying cellular and molecular mechanisms are distinct from each other. Although there is no single experimental model that precisely mimics human IBD, the localization of HSPCs in the MLN biopsies from IBD patients may at least suggest a common mechanism.

GM-CSF is an inflammatory cytokine produced by both hematopoietic (e.g., lymphocytes, innate lymphoid cells) and non-hematopoietic cells (e.g., fibroblasts, endothelial cells), and is required for HSPC proliferation and the differentiation and maturation of myeloid cells (Hamilton, 2019). We demonstrate that *Bacteroides* induces expansion of Ly6G^+^ myeloid cells in the spleen and MLN through innate immune signals, and Ly6G^+^ cells serve as tissue repairing cells that protect mice from inflammation-associated tissue damage during colitis (Figure 4F-G). Systemic injection of anti-Gr-1 antibody resulted in the depletion of both Ly6G^+^ (nearly 100%) and Ly6C^+^ (50%) cells (Fig. S4F), but as Ly-6C^+^ monocytes are inflammation-promoting cells that exacerbate colitis (Zigmond et al., 2012), we saw substantial expansion of both Ly6C^+^/G^+^ cells in DSS-treated MLN that expresses anti-inflammatory and immunosuppressive genes, and are reminiscent of MDSCs (Gabrilovich and Nagaraj, 2009). MDSCs are immunosuppressive and repress proliferation of antigen-specific T cells via nitric oxide secretion (Haile et al., 2008). Supportive of our findings, in T cell-mediated colitis CD11b^+^Gr-1^+^ cells expand and accumulate in the spleen and colon but not in the MLN through GM-CSF and IFN*γ* (Griseri et al., 2012), both known to be involved in MDSC generation (Ribechini et al., 2017). MDSC-related genes such as *Nos2*, *Cybb* and *Il6* (Veglia et al., 2018; Wu et al., 2012) were upregulated in MLN CD11b^+^Gr-1^+^ cells including both Ly6C^+^/G^+^ cells upon DSS treatment (Fig. 5K), as well as the anti-inflammatory M2 macrophages markers, *Cd163* and *Il4* (Mantovani et al., 2013) in Ly6C^+^/G^-^ cells.

Aging accompanies a decrease in intestinal barrier integrity, and thereby allows systemic microbial infiltration (Jeong et al., 2017). Here, both HSPC and Ly6G^+^ cells expanded in the BM and MLN of aged animals (Fig. 7D-F), and suggests the continuous stimuli from infiltrating microbiota as a possible driver for hematopoietic aging (Kovtonyuk et al., 2016). Upon depletion of the microbiota in germ-free animals, the aging-associated expansion of HSPC pool size (Beerman et al., 2010) was significantly suppressed (Fig. 7F). Whether microbiota deficiency under germ-free conditions can rescue or at least delay HSC aging in terms of reduced frequency and/or potential requires further functional experiments such as limiting dilution transplantations. A recent study with a cohort of ulcerative colitis (UC) patients demonstrated that an increased level of inflammatory cytokines enhances the risk for clonal hematopoiesis of indeterminate potential (CHIP) (Steensma et al., 2015), a pre-malignant condition associated with an aging-dependent accumulation of somatic mutations in HSCs (Zhang et al., 2019). Consistently, deficiency of Tet2, one of the key mutations for CHIP leads to a leaky gut and subsequent microbial translocation into the circulation, causing inflammation-promoting expansion of preleukemic stem cells in the mouse (Meisel et al., 2018). In contrast to acute colitis which affects the function of MPPs rather than HSCs, chronic gut inflammation compromises HSC transplantability and immunoreactivity against infection. Uncovering the mechanisms underlying gut-associated inflammation will help to understand feedback signals derived from the cross-organ communication that tailors hematopoiesis towards tissue surveillance and repair, and which might be relevant to aging-associated chronic disorders.

## Materials and methods

### Mice

C57BL/6 (CD45.2^+^) and B6.SJL (CD45.1^+^) mice and *Trif*^-/-^;*Myd88*^-/-^ mice (CD45.2^+^) were purchased from Japan SLC, Jackson Laboratories and Oriental BioService, respectively. *Hlf*^tdTomato+^ reporter and germ-free mice were kindly provided by Drs. Tomomasa Yokomizo (Kumamoto University) (Yokomizo et al., 2019) and Hiroshi Ohno (RIKEN). All mice were maintained at the Center for Animal Resources and Development at Kumamoto University. All experiments were approved by the Animal Care and Use Committee of Kumamoto University.

### Bacterial lysate preparation

The various strains of *Bacteroides* (*Bacteroides dorei*, *Bacteroide thetaiotaomicron*, *Bacteroides ovatus*, *Bacteroides stercoris*, *Bacteroides uniformis* and *Bacteroides vulgatus*) were purchased from RIKEN Bioresource Research Center. *Bacteroides* and *Feacalibacterium praunitzii* was grown in GAM medium (Nissui) under anaerobic conditions in Bactron 300 (Shellab), collected and centrifuged at 7,000 rpm for 5 minutes, 4°C and resuspended in PBS. The bacterial suspension was sonicated with a UD-100 sonicator (TOMY), maximum power with 30 second intervals and the lysate obtained by ultracentrifugation at 100,000 g for 60 minutes, 4°C (CS 100FNX, HITACHI) resuspended in PBS and the concentration was measured by Bio-Rad protein assay dye reagent concentrate (BioRad). Mice were *i.p.* injected with 15μg of the bacterial lysate or PBS, and analyzed by FACS 12 hours later. Three thousand MPP2 (LSKCD48^+^CD150^+^) cells of BM were sorted into 96 well plate containing 100μl of SF-O3 medium with 10% FBS and cultured with 100x GlutaMAX Supplement, 50x penicillin/streptomycin, 10ng/ml SCF, 10ng/ml IL-3, 10ng/ml GM-CSF, 50ng/ml EPO, 50ng/ml TPO, and 10mg/ml *Bacteroides* lysate. GM-CSF receptor expression levels were analyzed by FACS 2 and 4 days after culture.

### DSS-induced colitis

Mice were treated with double distilled water (ddW) with or without 2.5% dextran sulfate sodium (DSS) (35-50 kDa; MP Biomedicals, Tokyo, Japan) for 7 days to induce acute colitis, and then switched to ddW.

For chronic colitis induction, mice were treated with 2.0% DSS or ddW for one week, ddW for another week for 3 cycles, and ddW until analysis. Peripheral blood parameters were assessed with Celltacα MEK-6358 (NIHON KOHDEN COPARATION).

### FACS analysis

All antibodies were purchased from Thermo Fisher Scientific or Biolegend unless otherwise stated. For the mature cell analysis, pre-incubation with anti-CD16/32 antibody (93) to block Fc*γ*R was followed by staining with antibodies against B220 (RA3-6B2), CD3*ε* (145-2C11), F4/80 (BM8), Ly6G (1A8), Ly6C (HK1.4) and CD11b (M1/70). For early hematopoietic cell analysis, cells were incubated with the biotinylated antibodies against the lineage (Lin) markers: NK1.1 (PK136), CD11b (M1/70), Ter119 (Ter119), Gr-1 (RB6-8C5), CD4 (GK1.5), CD8*α* (53-6.7), CD3*ε* (145-2C11), B220 (RA3-6B2), and IL-7R*α* (SB/199)), and the fluorescence-conjugated antibodies: c-Kit (2B8), Sca-1 (D7), CD34 (HM34), Flt3 (A2F10), CD150 (TC15-12F12.2), CD48 (HM48-1), and CD16/32 (93). For common lymphoid progenitor (CLP) staining, IL-7R*α* (A7R34) was excluded from the Lin markers, and stained separately to define CLP. For multipotent progenitor (MPP) 2 sorting for RNA sequencing, total BM and mesenteric lymph node (MLN) cells were stained with Lin, c-Kit, Sca-1, CD48, CD150, and sorted on FACS Aria III (BD Biosciences). For donor chimerism analysis of mature hematopoietic cells, antibodies against CD45.1 (A20), CD45.2 (104), B220 (RA3-6B2), CD3*ε* (145-2C11), Gr-1 (RB6-8C5) and CD11b (M1/70) were used. The following antibodies were used for staining HSPC and mature populations of human MLN and BM samples: CD38 (HIT2), CD45RA (HI100), CD90 (5E10), CD19 (HIB19), CD16 (3G8), CD33 (WM53), CD14 (HCD14), CD3 (UCHT1), CD34 (581). FACS data was analyzed using FlowJo (BD).

### EdU incorporation

WT mice were *i.p.* injected with 100μg of EdU (Sigma Aldrich). BM and MLN cells were harvested and stained with antibodies against the Lin markers, followed by intracellular staining of EdU according to the instructions of Click-iT Plus EdU Alexa Fluor 488 Flow Cytometry Kit (Thermo Fisher, C10632). EdU positivity was defined by measuring the baseline intensity of non-EdU treated cells stained with the anti-EdU antibody.

### *In vitro* colony formation assay

For the CFU assay, 3x10^4^ BM or 3x10^6^ MLN cells were cultured in 1.5ml methylcellulose (StemCell Technologies, M3234) with standard cytokines: 10ng/ml SCF, 10ng/ml IL-3, and 10ng/ml GM-CSF. Colonies were classified by morphology after 14 days. For the single-cell colony assay, MPPs were sorted into a 96-well plate containing medium and 1mg/ml *Bacteroides* lysate and classified after 4-14 days. Three thousand BM MPP2 cells were sorted, cultured in standard medium with 10mg/ml *Bacteroides* lysate and the GM-CSFR expression was analyzed, 2 and 4 days later.

### ELISA

The BM fluid was flushed out using a 25G needle and syringe filled with PBS containing 0.1% BSA, centrifuged at 3,500 rpm for 5 minutes, 4°C and filtered with a 0.45μm filter (Merck, Germany). MLNs were mashed by sliding two glass slides together, centrifuged at 1,500 rpm for 5 minutes, 4°C and lysed with 500μm 1x RIPA buffer (Abcam, ab156034). PB was sampled with heparin-coated capillaries (HIRSCHMANN, Germany), and the serum was collected by centrifugation at 3,500 rpm for 10 minutes twice, 4°C. Microwell plates (BioLegend, #423501) were coated with purified GM-CSF capturing antibody from ELISA MAX™ Deluxe Set Mouse GM-CSF (Biolegend), left overnight at 4°C, and blocked with blocking buffer. Samples were added and incubated at room temperature (RT) for 2 hours, followed by detection of GM-CSF with HRP-conjugated detection antibody and TMB solution according to the manufacturer’s instructions (Biolegend).

### Antibiotics treatment

Four weeks-old mice were treated with 0.1% neomycin (NM) (Nacalai Tesque, Japan), 0.1% metronidazole (MNZ) (FUJIFILM Wako, Japan), 0.05% vancomycin (VCM) (FUJIFILM Wako, Japan), 0.1% ampicillin (ABPC) (FUJIFILM Wako, Japan), or a mixture of all four in the drinking water for 4 to 8 weeks. Mice were subsequently treated with DSS or ddW for another week, and the phenotype was analyzed by FACS. Kaplan-Meier survival was also analyzed. The genomic DNA of gut microbiota was extracted from murine cecal contents, and 16S rRNA gene sequencing was conducted using a MiSeq sequencer as previously described (Ishii et al., 2018). The microbiome analysis data have been deposited in the DNA Data Bank of Japan (DDBJ) Sequence Read Archive (http://trace.ddbj.nig.ac.jp/dra/) under the accession number DRA009488.

### Ly6C^+^/G^+^ cell depletion

DSS treated mice were *i.p.* injected with 500μg of anti-Gr-1 (Ly6C/G) neutralizing antibody (RB6-8C5) (BioXCell) or IgG2b isotype-matched control (BioXCell) 6 days after the start of DSS treatment. Mice were sacrificed at day 8 and the BM, MLN, and PB were analyzed. The clinical disease severity was monitored daily and scored as previously described (Wirtz et al., 2017): a) weight loss, score 0 = 0%; score 1 = 1-5%, score 2 = 6-10%, score 3 = 11–18%, score 4 = >18%; b) stool consistency, score 0 = normal, score 1 = soft but still formed, score 2 = soft, score 3 = very soft or wet, score 4 = watery diarrhea; c) rectal bleeding, score 0 = negative hemoccult and normal stool, score 1= negative hemoccult but abnormal stool, score 2 = positive hemoccult, score 3 = traces of blood visible in stool, score 4 = gross rectal bleeding. For hematoxylin and eosin (H&E) staining, the entire colon was dissected from measured, and the middle part of the colon sampled. Colon tissues were fixed with 10% formaldehyde, overnight at 4°C and paraffin embedded, sectioned, and H&E stained. The histopathological scores of H&E stained samples were examined in a double-blinded manner according to the scores below:

1. Disruption of intestinal epithelium structures: score 1= 0-25%; score 2= 25-50%; score 3= >50%
2. Formation of ectopic follicles: score 0= not formed; score 1= ectopic follicles formed
3. Loss of goblet cells: score 1= 0-25%; score 2= 25-50%; score 3= >50%
4. Infiltration of immune cells: score 1= 0-25%; score 2= 25-50%; score 3= >50%
5. Attachment and/or invasion of bacteria into intestinal epithelium: score 0= none; score 1= bacteria attachment/invasion of bacteria detected.

Final scores were calculated by the Student’s t-Test (two-tailed t-test).

### Serial BM transplantation

Total BM cells were isolated from chronic DSS- or ddW-treated CD45.2^+^ mice at 2-3 months post treatment, and *i.v.* transplanted into lethally-irradiated (10Gy) CD45.1^+^ mice. PB donor chimerism was assessed every 4-6 weeks up to 24 weeks. Mice were sacrificed to check for BM donor chimerism. For secondary transplantation, 2x10^6^ total BM cells from 1^st^ recipients were transplanted into lethally irradiated CD45.1^+^ mice. PB and BM donor chimerism were similarly assessed. Kaplan-Meier survival was analyzed.

### LPS challenge

WT mice were exposed to chronic DSS treatment with 3 cycles of one week ddW/DSS followed by one week ddW. Four weeks after final DSS treatment, mice were *i.p.* injected with 100μg of LPS or PBS, and BMs harvested 16 hours post injection were analyzed using FACS Aria III.

### PTX challenge

DSS treated mice were *i.v.* injected with 1μg of pertussis toxin (PTX) (Enzo, BML-G100-0050) or PBS at day 4 of 1.5% DSS treatment, sacrificed at day 7 and the BM and MLN were analyzed by FACS. The BW change and survival rate in each group were monitored daily. For H&E staining, the entire colon was dissected from each mouse and followed by sampling of the middle part of the colon. Colon tissues were fixed with 10% formaldehyde, overnight at 4°C and paraffin embedded, sectioned, and H&E stained. The histopathological scores of H&E stained samples were examined in a double-blinded manner and final scores were calculated by the Student’s t-Test (two-tailed t-test).

### Lymph node imaging

MLN harvested from DSS-treated *Hlf*^tdTomato+^ mice at day (d)7 were fixed overnight in 4% PFA at 4°C, washed and cryoprotected with 30% sucrose in PBS overnight at 4°C. Samples were stained and cleared using the iDISCO method as previously reported (Renier et al., 2014). Briefly, MLN samples were washed in PTx.2 (0.2% TritonX-100 in PBS), incubated in pretreatment solution 1 (0.2% TritonX-100 and 20% DMSO in PBS) overnight, and pretreatment solution 2 (0.1% Tween-20, 0.1% TritonX-100, 0.1% Deoxycholate, 0.1% NP-40 and 20% DMSO in PBS) for 7-8 hours. After washing in PTx.2, samples were treated with the permeabilization solution (0.3M Glycine and 20% DMSO in PBS) overnight at 37°C, followed by the blocking solution (6% goat serum and 10% DMSO in PTx.2) for 7-8 hours at 37°C. Soutamples were stained in primary antibody solution (5% DMSO, 3% goat serum in PTwH containing 0.2% Tween-20, heparin in PBS) with primary antibodies (RFP, CD31 and PNAd) for 1.5 days at 37°C, washed overnight with PTwH and stained in the secondary antibody solution (3% goat serum in PTwH) with secondary antibodies (donkey anti-Rabbit IgG (H+L) conjugated with Cy-3, goat anti-Armenian Hamster IgG (H+L) conjugated with Alexa Fluor 488 and donkey anti-Rat IgG (H+L) conjugated with Alexa Fluor 647) at 37°C for 6 hours. Samples were washed in PTwH overnight and dehydrated with a stepwise increase of methanol (20%, 40%, 60%, 80%, and 100% methanol in ddW), 66% Dichloromethane (DCM) in methanol, 100% DCM, and cleared with di-benzylether until transparent. Of note, all incubation and washing steps were done on a shaker/rotar, at room temperature (RT) unless otherwise stated. Samples were imaged with the Leica SP8 confocal microscope. Three-dimensional (3D) reconstructions were generated from z-stack images in Imaris (Bitplane). Distance between tdTomato^+^ cells to the nearest vasculature were calculated by digital spots and surface creation with the Distance Transformation Matlab XTension package on Imaris. A separate channel of randomly generated spots was created in ImageJ using the plugin called 3D random ellipsoids downloaded online (https://imagejdocu.tudor.lu/doku.php?id=plugin:stacks:3d_tools:start) (Ollion et al., 2013). Of note, when using this macro, dimensions of the channel were set to be identical to the 3D image (SizeX, SizeY, SizeZ), and the radius X and Y were set to the same value. The resulting channel was saved as a tiff file and overlaid onto the original 3D reconstructed image.

### Quantitative RT-PCR

Cells were sorted directly into a 96-well plate containing cell lysis buffer (Roche), and frozen immediately until random displacement amplification (RamDA) as previously reported (Hayashi et al., 2018). Briefly, the first strand of cDNA was synthesized with the PrimeScript RT reagent kit (TAKARA Bio Inc.) and NSR (not-so random) primers. The second strand was synthesized with Klenow Fragments (3′-5′ exo-; New England Biolabs Inc.) and complement chains of NSR primers. Quantitative PCR was performed using SYBER green (Applied Biosystems) on a Real-time PCR LightCycler96 (Roche). The relative mRNA expression of each gene was normalized against *Gapdh* and *B2m* expression. Refer to supplemental table for primer sequences.

### RNA sequencing

A hundred MPP2 cells were sorted from the BM and MLN, and 100 HSC^LT^ cells were sorted from the BM. Samples were subjected to the RamDA method as described in Quantitative RT-PCR. Following second strand synthesis, the double-stranded cDNA was purified with AMPure XP beads (Beckman Coulter, Inc. CA), and subjected to library preparation using the Nextera XT DNA sample Prep kit (Illumina Inc.). The quantity and quality of the isolated cDNA library was determined with Multina (SHIMADZU CORPORATION, Kyoto, JAPAN) and Bioanalyzer 2100 (Agilent, Waldbronn, Germany), and the library was sequenced on the Next-Seq system (Illumina Inc.). Quality check and trimming of single-end sequences were completed with the trim_galore (version 0.4.3) package. Quality and length parameters were --quality 30 and --length 30. Filtered sequences were processed to remove mitochondrial mRNA with the SortMeRNA and were aligned to mouse reference sequences (GRCm38, M17 version) from (http://www.genecodesgenes.org/mouse/release.html) with ultrafast RNA-seq aligner STAR (version 2.5.3a) (Dobin et al., 2012). All aligned bam files were used with mouse GFF annotation files (GRCm38, M17 version) as input into the featureCounts program from the Subreads program (version 1.6.1) to count the raw reads for each gene and sample, and to create a gene count matrix for MLN and BM samples. To calculate differentially expressed genes, the DESeq2 (version 1.20.0) package was used in R (version 3.4). For genes to be considered significantly and differentially expressed, alpha = 0.05 was used.

### Patient and tissue samples

Human bone marrow (hBM) aspirates were obtained from patients requiring a total knee or hip arthroplasty, while human MLN samples were obtained from patients with inflammatory bowel disease (IBD) requiring intestinal resection at Kumamoto University Hospital. The study was approved by the Medical Ethics Committee of Kumamoto University (Approval No. 960 and IRB approved number: 1725, respectively), and all patients signed the informed consent form to participate.

### Quantification and statistical analysis

All data are shown as the mean ± s.e.m., unless otherwise indicated. Kaplan-Meier survival was analyzed using BellCurve for Excel (Social Survey Research Information Co., Ltd.). All other statistics were performed with the Student’s t-Test (two-tailed t-test): ns, not significant; \**p*<0.05; \*\**p*<0.01; \*\*\**p*<0.001.

## Online supplemental material

Fig. S1 contains hematopoietic changes induced by acute colitis upon DSS treatment in mouse PB, BM, spleen and PLN (A-G) as well as HSPCs and mature populations found in the MLN of human IBD samples and human BM (different donors) as a staining control (H-K). Fig. S2 includes hematopoietic changes induced by DSS in WT and *Trif*;*Myd88* DKO mice, and suggests that innate immune signals control DSS-induced acute colitis. Fig. S3 contains results of various antibiotics treatment for narrowing down of the microbiota responsible for the immature and mature hematopoietic changes induced upon acute colitis, and *in vitro* CFU data of MPPs when cultured with GM-CSF and *Bacterioides*. Fig. S4 shows kinetics of Ly6C^+^/G^+^ in the BM and PB over 4 to 12 days upon DSS treatment and results of their depletion using the Gr-1 neutralizing antibody. Fig. S5 includes changes in blood parameters and HSC function tested by transplantation experiments induced by chronic colitis.

## Acknowledgments

We would like to thank the International Core-facility of Advanced Life Science at Kumamoto University and Ms. Sayuri Nakata for their technical assistance, Ms. Ryoko Koitabashi for preparation of the RNA-Seq library, Dr. Akifumi Kiyota for mass cytometric analysis, and Mr. Tomoya Tsukimi for microbiome data deposition. This work was supported by KAKENHI from the Japanese Society of the Promotion of Science (JSPS) (18K16091 to Y.H., 17H05654 and 18H04805 to S.F., 15H01519 and 17H05651 to H.T.), KAKETSUKEN and MEBAE research programs at Kumamoto University (to Y.H.), JSPS fellowship (201820690) and a grant for Excellent Graduate Student at Kumamoto University (to M.S.), JST PRESTO (JPMJPR1537 to S.F.), JST ERATO (JPMJER1902 to S.F.), AMED-CREST (JP19gm1010009 to S.F.), the Takeda Science Foundation (to S.F.), the Food Science Institute Foundation (to S.F.), the Program for the Advancement of Research in Core Projects under Keio University’s Longevity Initiative (to S.F.), KANAE Foundation for the Promotion of Medical Science (to H.T.), SENSHIN Medical Research Foundation (to H.T.), Astellas Foundation for Research on Metabolic Disorders, Mochida Memorial Foundation (to H.T.), Princess Takamatsu Cancer Research Fund (to H.T.), MIRAI Research program and Center for Metabolic Regulation of Healthy Aging at Kumamoto University (to H.T.).

## Authors’ contribution

Y.H. and M.S. designed and performed the experiments, analyzed the data, and wrote the manuscript. T.M., M.F., J.M., S.A., P.K. G.N. and S.F. performed the experiments and the analyzed data. S.B analyzed the RNA sequencing data. Y.M. and H.B. provided human biopsies via IRB approved by Kumamoto University. H.T. designed the experiments, supervised the research project and wrote the manuscript. Y.H. and M.S. contributed equally to this study.

## Figure Legends

**Figure S1 DSS treatment induces acute colitis and hematopoietic changes.** (A) Time-course kinetics of PB parameters (RBC, red blood cell; WBC, white blood cell; Hb, hemoglobin; Plt, platelet) and changes in body weight (BW). PB was taken every 3-4 days after DSS treatment for analysis (n=5-9 from 2 experiments). (B) Representative FACS plots of HSPCs in the BM from ddW- or DSS-treated mice at day (d)8. (C) Representative FACS plots of HSPCs in the PB of ddW- or DSS-treated mice. (D) Absolute number (upper) and proportion (lower) of HSPCs in the PB of mice treated with ddW (blue) or DSS (red) at d5, d8, and d10 (n=3-5 from 2 experiments). (E) Absolute number of HSPC subpopulations in the spleen of ddW (blue)- or DSS (red)-treated mice at d8 (n=11 from 4 independent experiments). (F) Representative FACS plots of HSPCs in the PLN of ddW- or DSS-treated mice. (G) Absolute number of HSPCs in the BM of mice treated with ddW (blue) or DSS (red) at d5, d8, and d10 (n=3-4 from 2 experiments). (H-I) Representative FACS plots (H) and absolute number (I) of HSPC populations in human MLN (hMLN) sampled from proximally and distally inflamed regions obtained from IBD patients and human BM (hBM) as a staining control (n=5 from 5 experiments). (J-K) Representative FACS plots (J) and absolute number (K) of mature populations in hMLN sampled from proximally and distally inflamed regions (n=6 from 6 experiments).

**Figure S2 Innate immune signals control immature and mature hematopoiesis.** (A) Absolute number of HSPCs in the spleen of WT or *Trif*;*Myd88* DKO mice at d8 upon acute colitis induction. WT-ddW (blue), WT-DSS (red), DKO-ddW (green), DKO-DSS (orange) (n=3-7 from 3 experiments). (B) Representative FACS plots of mature cell populations in the BM of ddW- or DSS-treated WT or DKO mice. (C-D) Absolute number of mature cell populations, B220^+^ B cells, CD3*ε*^+^ T cells, Ly6C/G^high^ cells or Ly6C/G^low^ cells in the BM (C) and spleen (D) of WT-ddW (blue), WT-DSS (red), DKO-ddW (green), and DKO-DSS (orange) at d8 upon colitis induction (n=3-7 from 3 experiments). \**p*<0.05;\*\**p*<0.01; \*\*\**p*<0.001 (two-tailed t-test).

**Figure S3 Effect of antibiotics, Bacteroides and innate immune signals on immature and mature hematopoietic changes.** (A) Unweighted (left) and weighted (right) UniFrac analysis of gut microbiota from antibiotics-treated mice. (B) Survival rate of antibiotics pretreated mice post DSS treatment (n=5 from 2 experiments). **p<0.01 (Kaplan-Meier estimate). (C-F) Absolute number of HSPCs in the BM (C), mature cells in the BM (D), spleen (E) and PB (F) of WT or *Trif*;*Myd88* DKO mice injected with PBS (blue/green) or *Bacteroides* (red/yellow) (n=3-4 from 3 experiments). (G) Experimental scheme for *in vitro* culture: 3000 MPP2 (LSKCD48^+^CD150^+^) cells were sorted from the BM of WT mice and analyzed by FACS 2 days after culture. (H) Representative histogram (left) and mean of intensity (right) of GM-CSFR*β* expression of BM MPP2 post *Bacteroides* stimulation *in vitro* (n=3 from 2 experiments). n.d., not detected; \**p*<0.05 (two-tailed t-test). (I) Experimental scheme for *in vitro* single-cell colony formation assay. (J) Time-course kinetics of colony-forming efficiency was evaluated every 2-3 days during culture (data from 3-4 experiments). (K-L) Colony-forming efficiency of BM MPPs (K) and MPP2 and MPP3/4 (L), 14 days after culture with or without addition of GM-CSF and *Bacteroides*. *,^$,¥,&,#^*p*<0.05; **,^$$,¥¥,##^*p*<0.01, n.s.; not significant (two-tailed t-test, *control vs GM-CSF alone, ^$^control vs *Bacteroides* alone, ^¥^control vs *Bacteroides* and GM-CSF; *^&^*GM-CSF alone vs *Bacteroides* alone, *^#^*Bacteroides alone vs *Bacteroides* and GM-CSF).

**Figure S4 Depletion of Ly6C+/G+ cells exacerbates colitis pathogenesis.** (A-C) Representative FACS plots (A), time-course kinetics of mature cell populations in BM (B) and PB (C) (n=3-4 from 2 experiments). B220^+^ B cells, CD3*ε*^+^ T cells, Ly6G^+^ cells, F4/80^+^Ly6C^+^SSC-H^low^ or F4/80^+^Ly6C^-^SSC-H^low^ monocytes and F4/80^+^SSC-H^high^ eosinophils. \**p*<0.05; \*\**p*<0.01; \*\*\**p*<0.001 (two-tailed t-test). (D) Disease activity index for BW loss, bleeding, and stool consistency post injection of neutralizing anti-Gr-1 antibody (*α*Gr-1) or IgG isotype-matched control (IgG) (n=4-7 from 3 experiments). (E) Absolute number of Ly6G^+^ and Ly6C^+^ cells in the BM and MLN of DSS treated mice after treatment with *α*Gr-1 or IgG (n=7-8 from 3 experiments). \**p*<0.05; \*\**p*<0.01;, \*\*\**p*<0.001. (F) Absolute number of HSPCs in the BM (left) and MLN (right) (n=4-8 from 3 experiments). \**p*<0.05; \*\**p*<0.01;, \*\*\**p*<0.001; ^#^*p*<0.05; ^##^*p*<0.01; ^###^*p*<0.001 (two-tailed t-test, *ddW-IgG compared to DSS-IgG; ^#^DSS-IgG compared to DSS-*α*Gr-1).

**Figure S5 Chronic colitis affects blood parameters and HSC function through epigenetic alterations.** (A) Time-course kinetics of PB parameters (RBC, red blood cell; WBC, white blood cell; Hb, hemoglobin; Plt, platelet) and BW in chronic colitis (ddW, blue; DSS, red). (B) Donor chimerism in overall, myeloid cells, B, and T cells in the BM of 1^st^ recipients (n=8-12 from 2 independent experiments). (C) Survival of 2^nd^ recipients post BMT (n=5 from 1 experiment). *p<0.05 (Kaplan-Meier method). (D) Time-course kinetics of donor chimerism in the PB of 2^nd^ recipients (n=5 from 1 experiment). (E-F) Donor chimerism in mature cells (E) and HSPCs (F) of BM post 2^nd^ BMT (n=5 from 1 experiment). (G) Experimental scheme of LPS re-challenge post chronic colitis: 4 weeks after chronic DSS treatment, mice were *i.p.* injected with 100μg of LPS or PBS and analyzed 16 hours later. (H) Absolute number of HSPCs in the BM upon LPS re-challenge (n=4-7 from 4 experiments).

## Tables

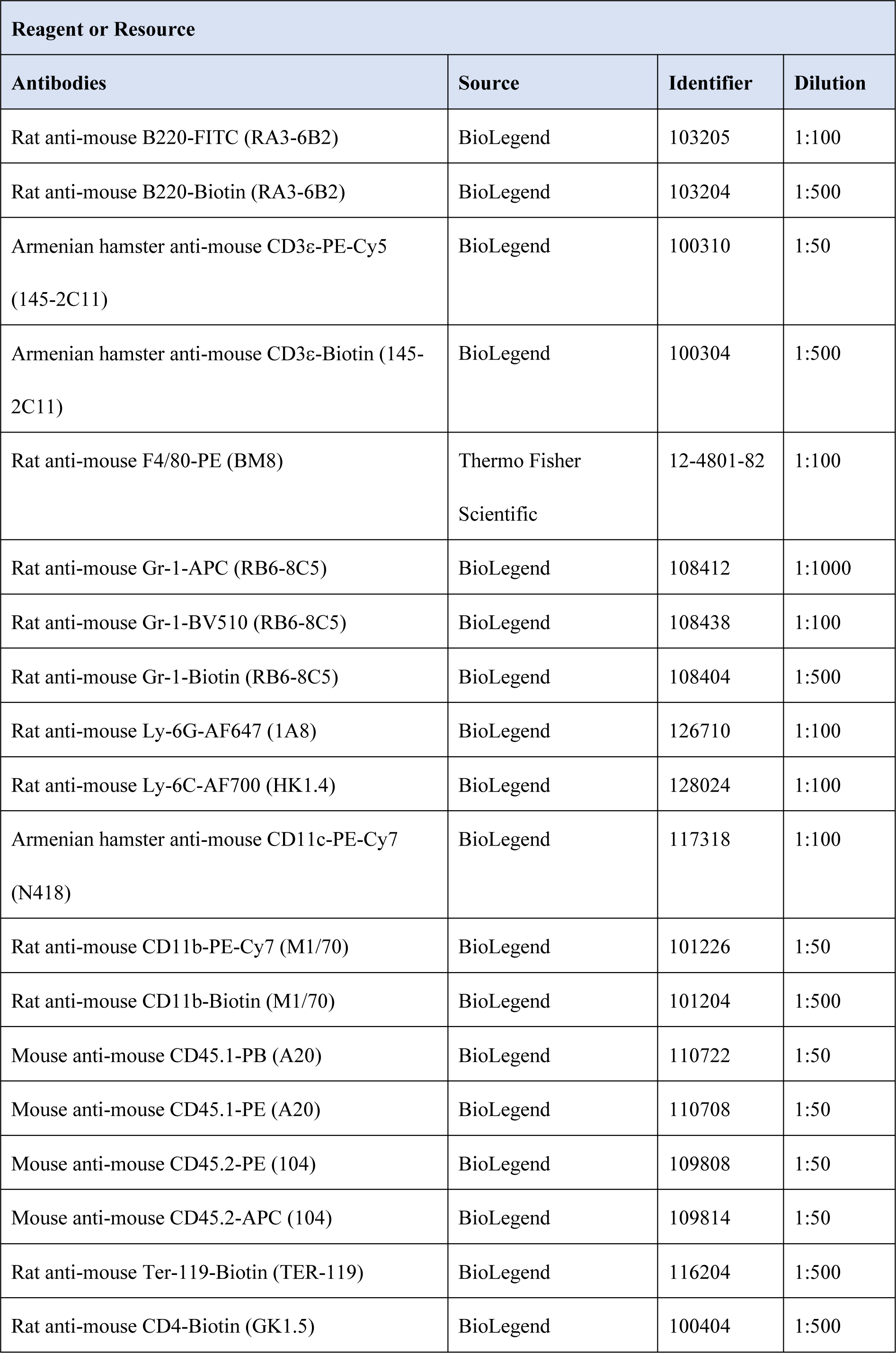
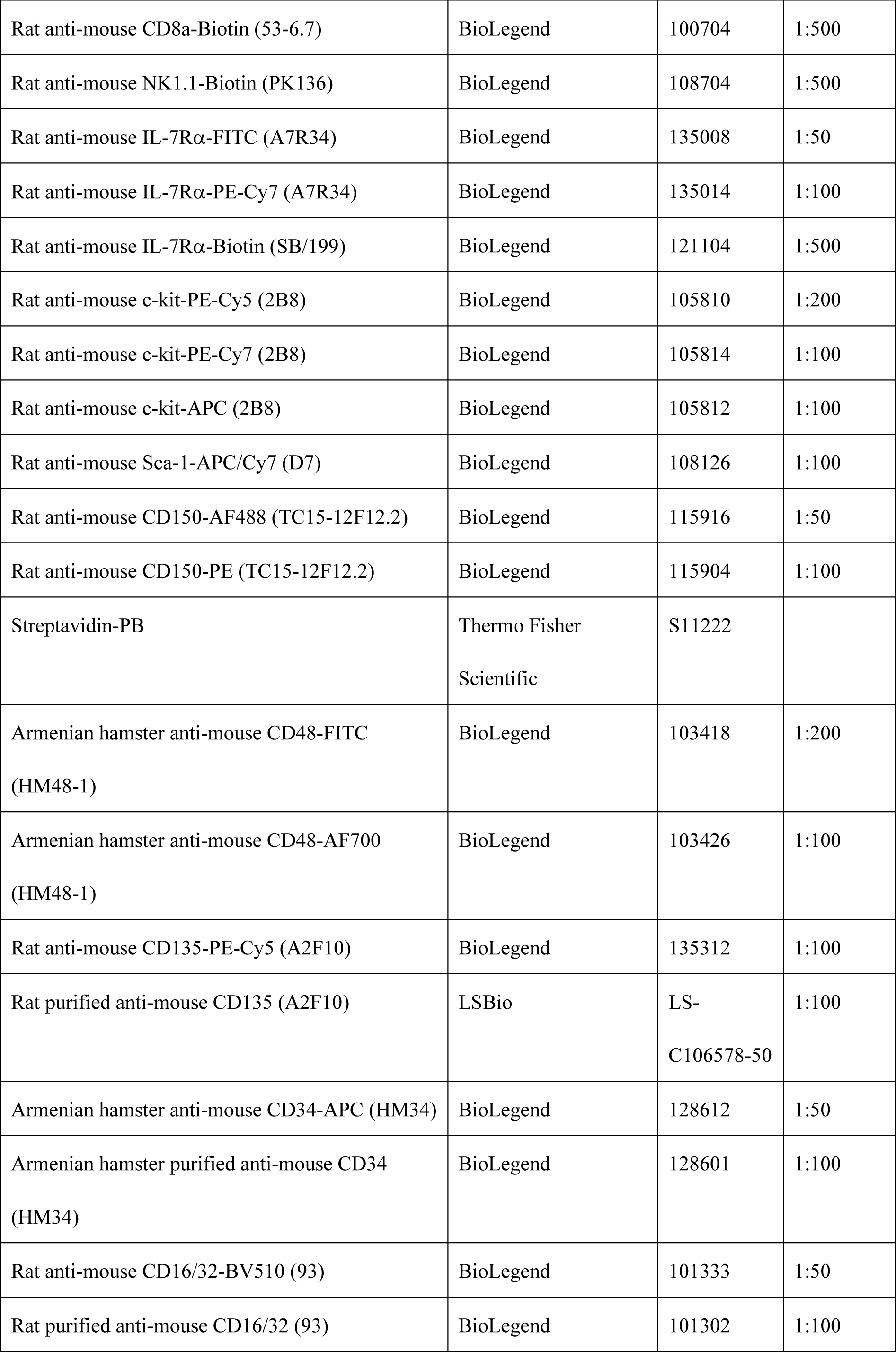
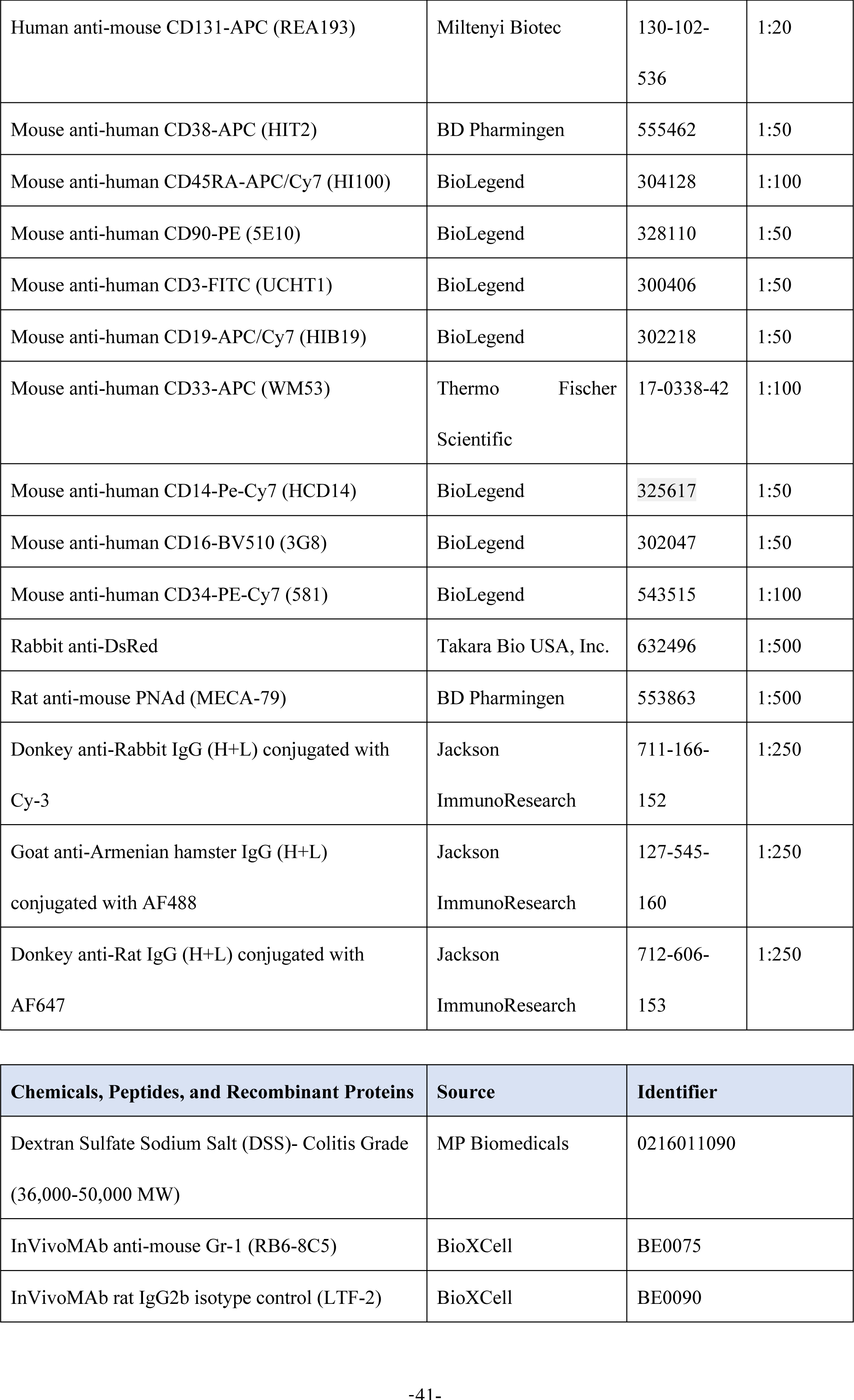
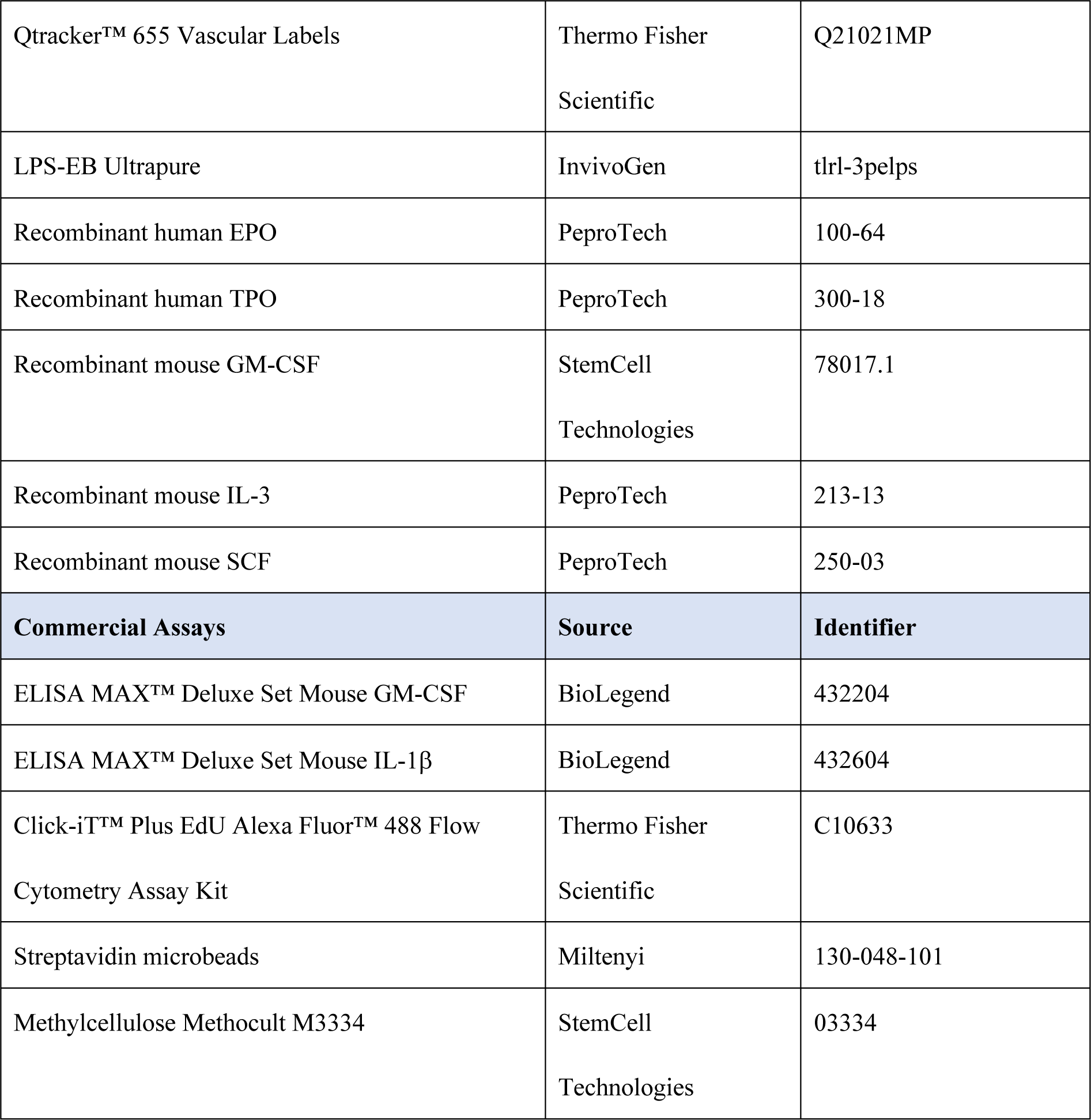
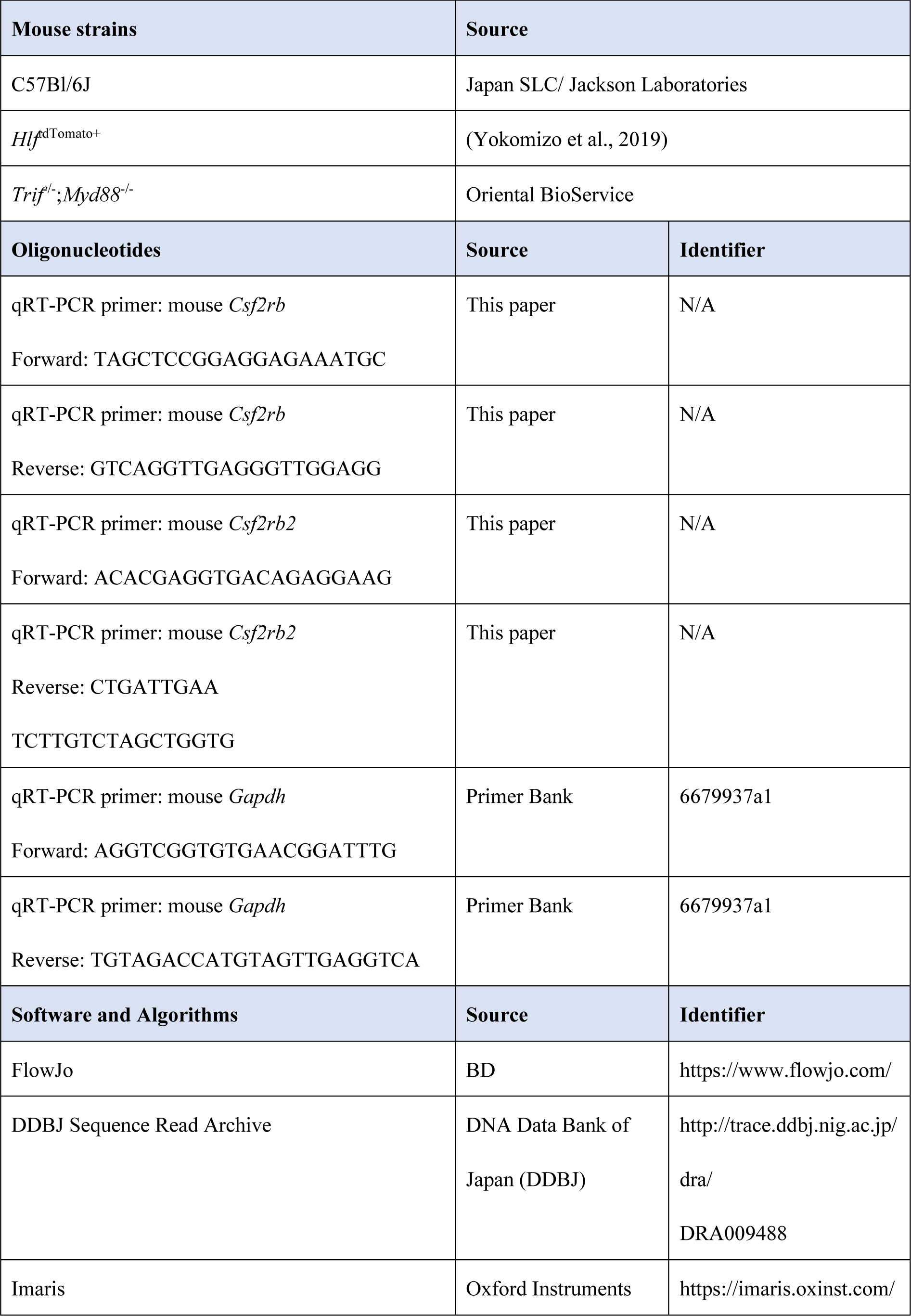
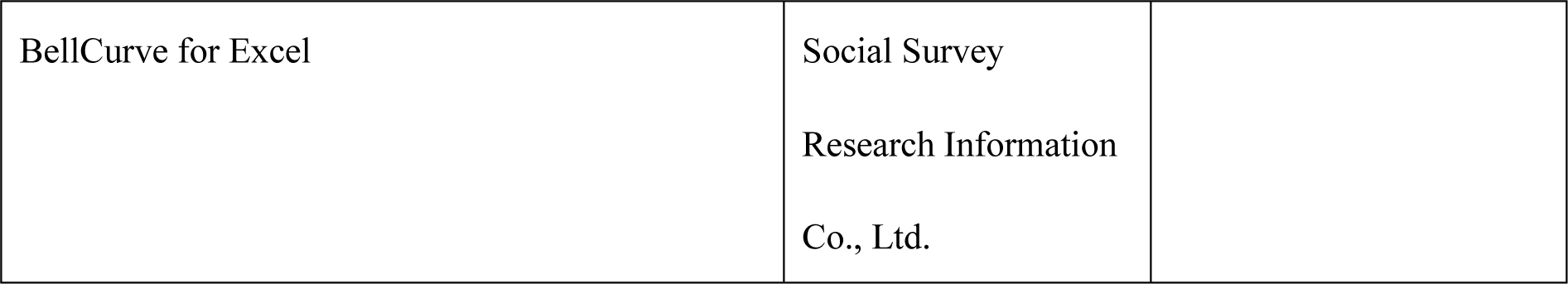

