## Supplemental figures for "Hematopoietic stem and progenitor cells integrate *Bacteroides*-derived innate immune signals to promote gut tissue repair"

### Figure S1 Hayashi et al.

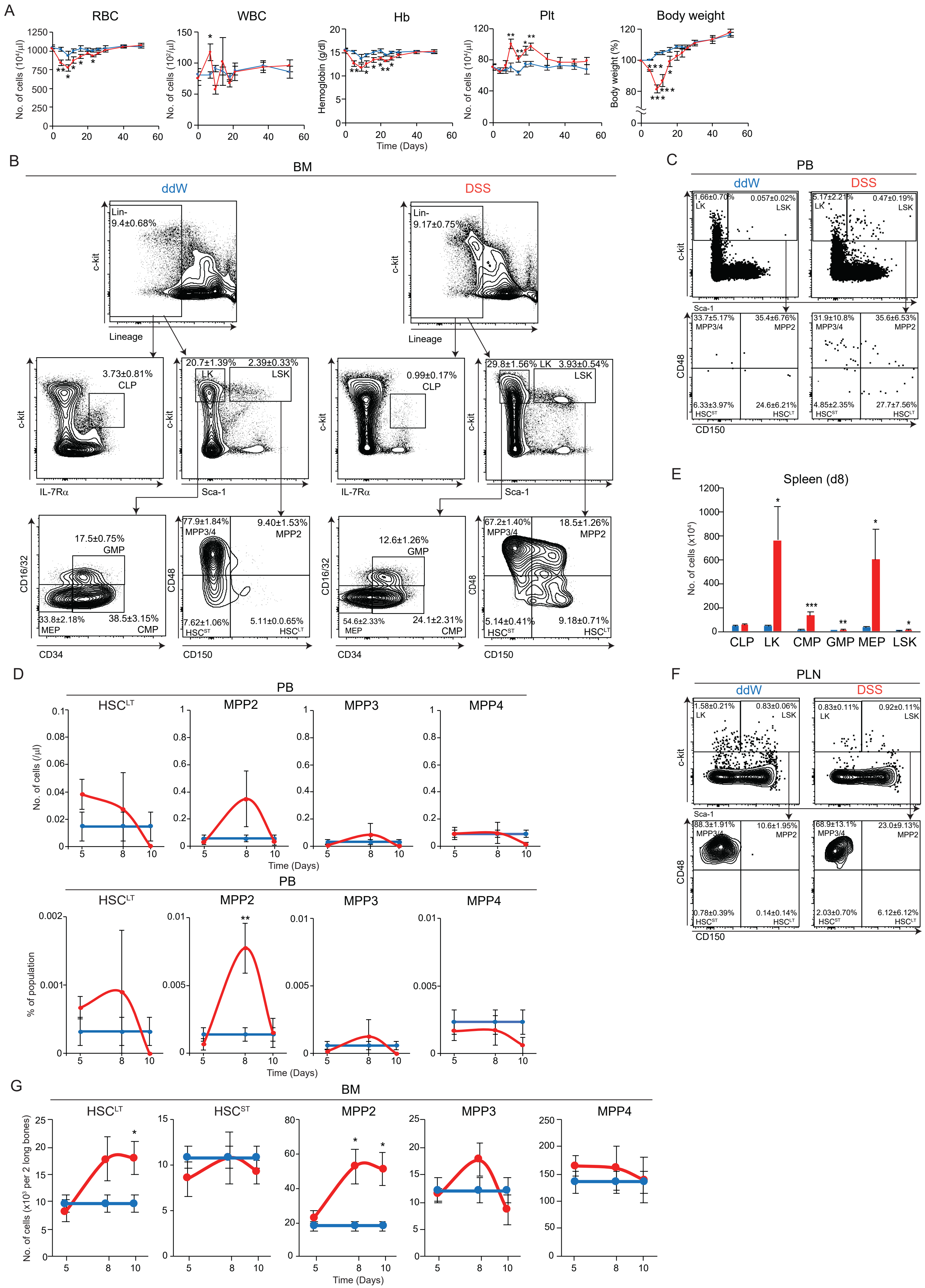

### Fig. S1 Hayashi et al. (Continued)

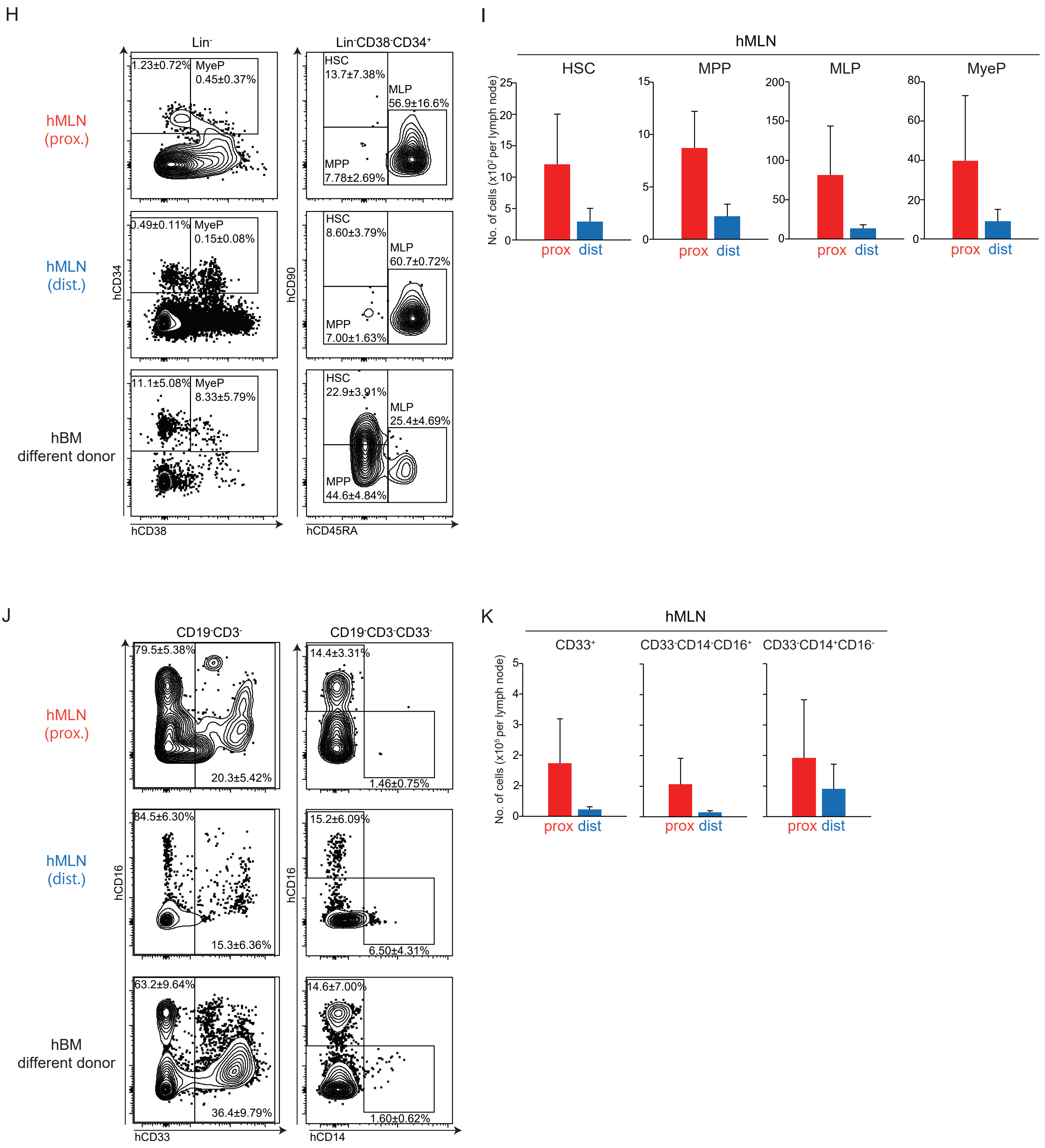

Figure S2 Hayashi et al.

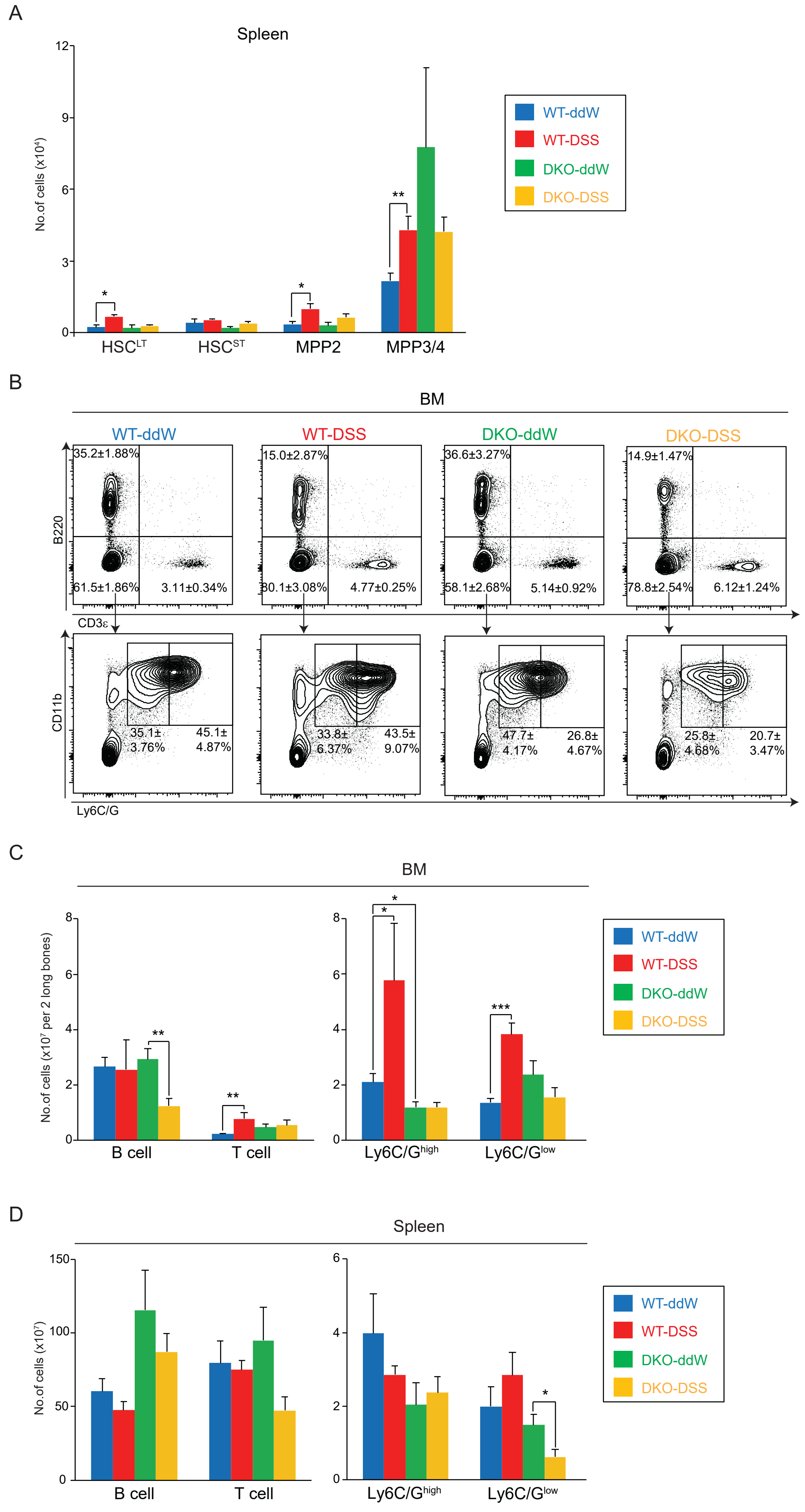

### Figure S3 Hayashi et al.

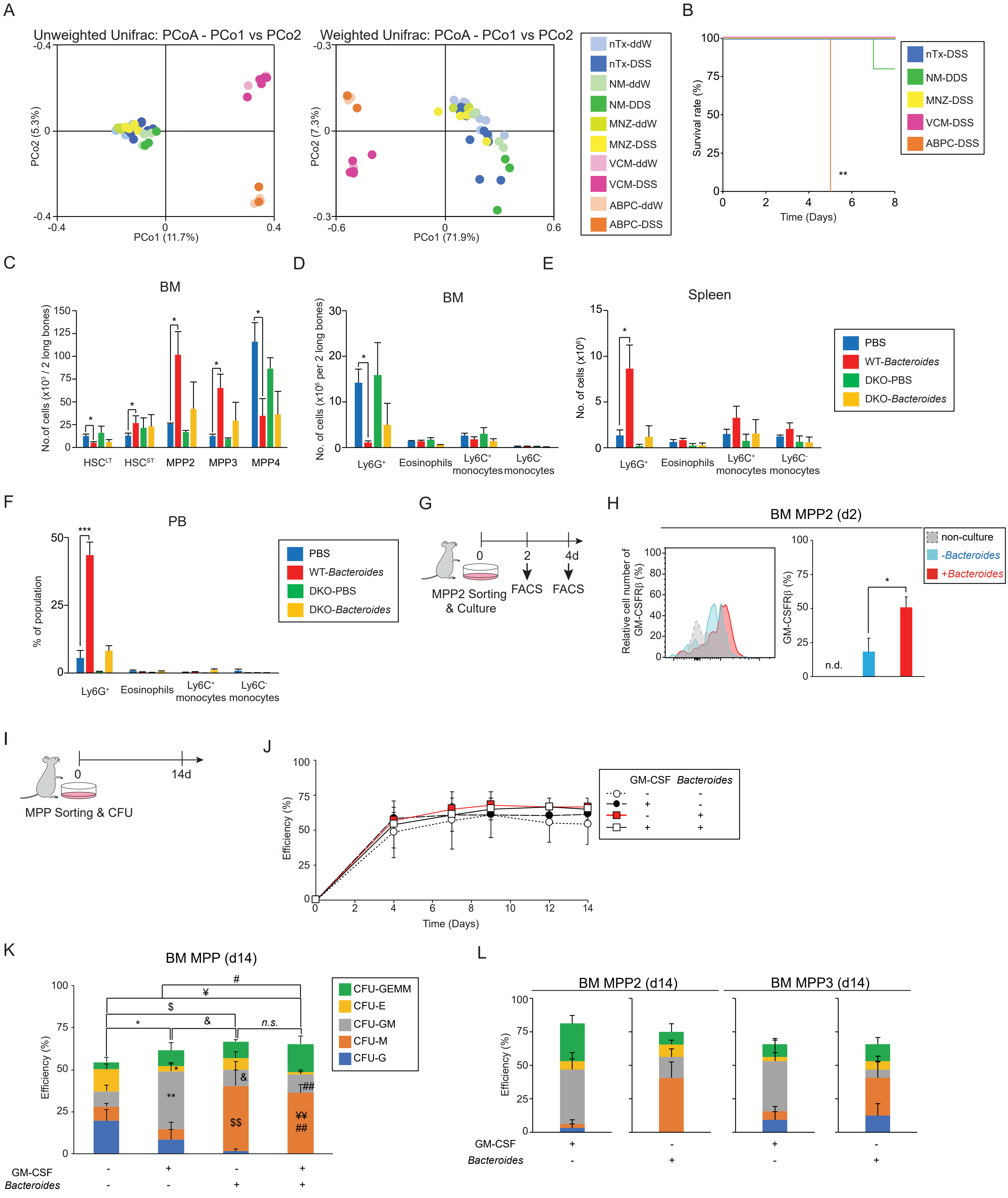

### Figure S4 Hayashi et al.

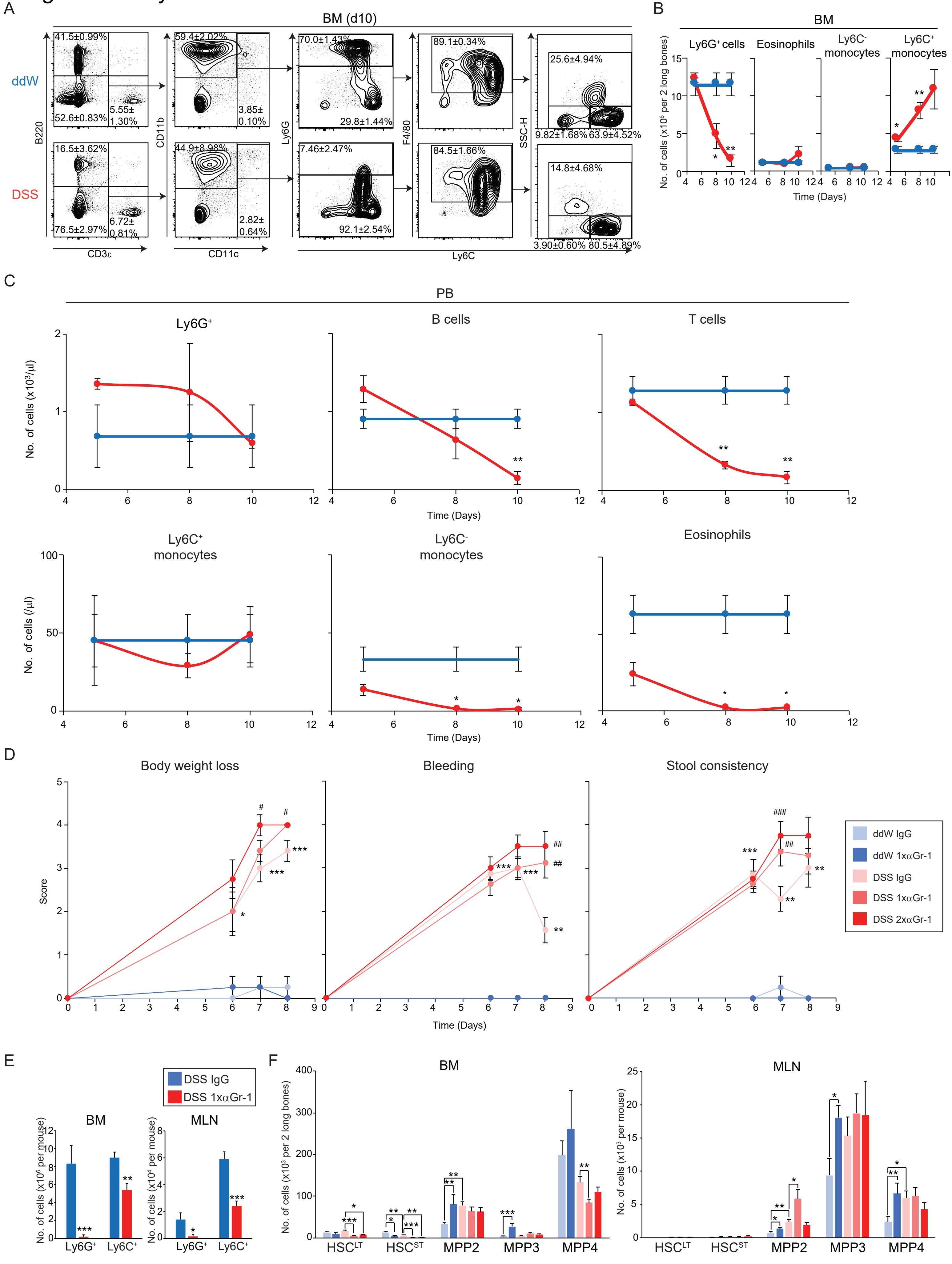

### Figure S5 Hayashi et al.

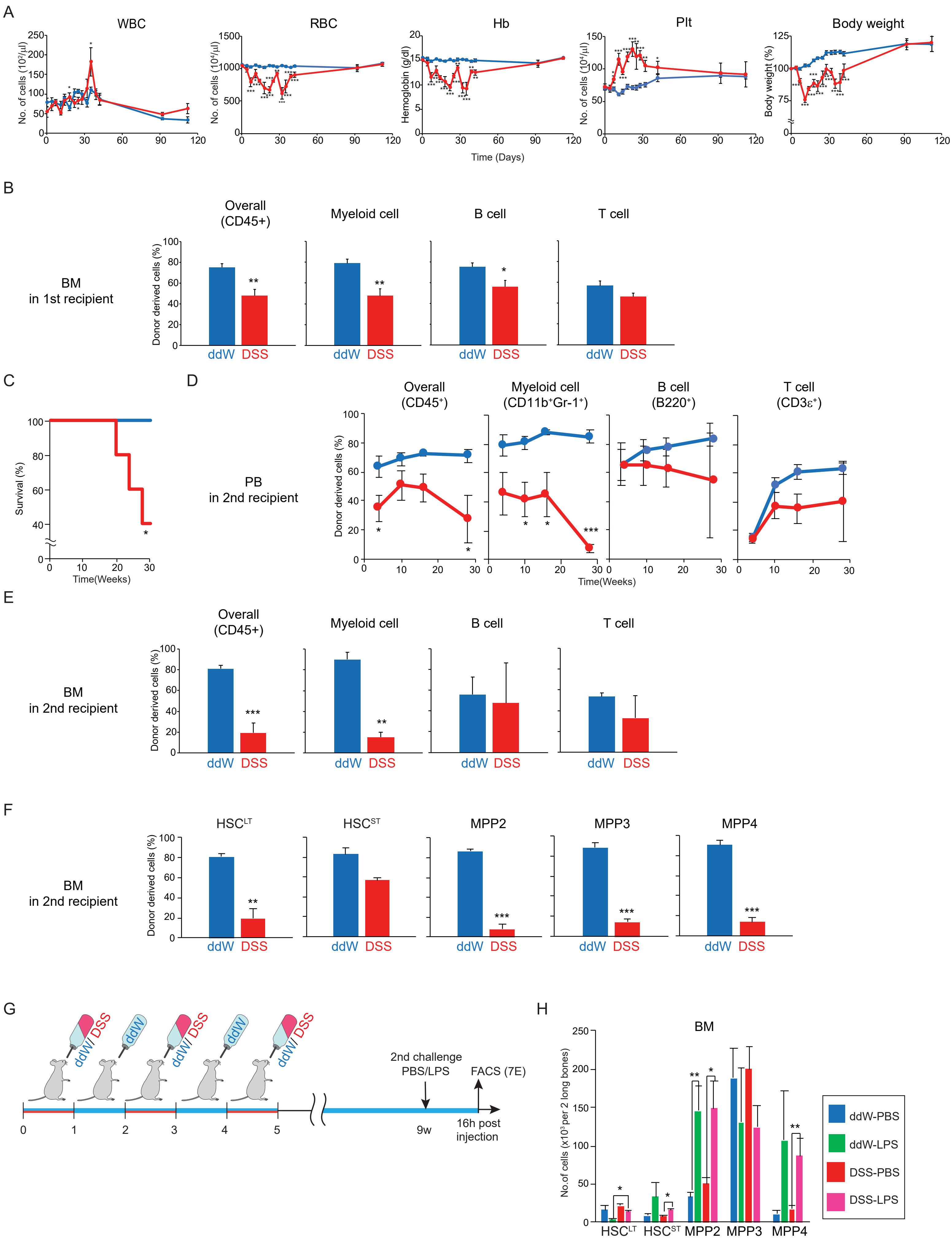
